# Novel step selection analyses on energy landscapes reveal how linear features alter migrations of soaring birds

**DOI:** 10.1101/805374

**Authors:** Joseph M. Eisaguirre, Travis L. Booms, Christopher P. Barger, Stephen B. Lewis, Greg A. Breed

## Abstract

Human modification of landscapes includes extensive addition of linear features, such as roads and transmission lines. These can alter animal movement and space use and affect the intensity of interactions among species, including predation and competition. Effects of linear features on animal movement have seen relatively little research in avian systems, despite ample evidence of their effects in mammalian systems and that some types of linear features, including both roads and transmission lines, are substantial sources of mortality. Here, we used satellite telemetry combined with step-selection functions designed to explicitly incorporate the energy landscape (el-SSFs) to investigate the effects of linear features and habitat on movements and space use of a large soaring bird, the golden eagle *Aquila chrysaetos*, during migration. Our sample consisted of 32 adult eagles tracked for 45 spring and 39 fall migrations from 2014-2017. Fitted el-SSFs indicated eagles had a strong general preference for south-facing slopes, where thermal uplift develops predictably, and that these areas are likely important aspects of migratory pathways. el-SSFs also revealed that roads and railroads affected movement during both spring and fall migrations, but eagles selected areas near roads to a greater degree in spring compared to fall and at higher latitudes compared to lower latitudes. During spring, time spent near linear features often occurred during slower-paced or stopover movements, perhaps in part to access carrion produced by vehicle collisions. Regardless of the behavioral mechanism of selection, use of these features could expose eagles and other soaring species to elevated risk via collision with vehicles and/or transmission lines. Linear features have been previously documented to affect the ecology of terrestrial species (e.g., large mammals) by modifying individuals’ movement patterns; our work shows these effects on movement extend to avian taxa.

## Introduction

Linear features, such as roads, railroads, and transmission line corridors, are major anthropogenic modifications to landscapes worldwide and disrupt the natural spatial heterogeneity of habitats. Linear features have changed how animals move (James and Stuart-Smith, 2000; Dyer et al., 2002; Whittington et al., 2004, 2005, 2011; Dickson et al., 2005; Latham et al., 2011; Dickie et al., 2017; Scrafford et al., 2018), which in turn has altered predator functional responses (McKenzie et al., 2012), increased stress levels in free-living animals (Wasser et al., 2011), and changed other ecosystem interactions (Haddad et al., 2003).

These effects arise via a myriad of mechanisms. Linear features change the permeability of the landscape (Dyer et al., 2002; Whittington et al., 2004; Dickson et al., 2005; McKenzie et al., 2012; Tremblay and Clair, 2009), the distribution of food (Latham et al., 2011; Whittington et al., 2011; McKenzie et al., 2012; Dickie et al., 2017), and the spatial distribution of mortality risk (James and Stuart-Smith, 2000; Latham et al., 2011; Whittington et al., 2011; DeGregorio et al., 2014; Popp et al., 2018). Some linear features have negative effects that are unidirectional in predator-prey interactions. Seismic lines, for example, increase a predator’s (wolf) access to prey (caribou), negatively affecting prey but imposing no additional risk or harm to the predator (McKenzie et al., 2012; DeMars et al., 2016; Dickie et al., 2019).

Many linear features, such as roads and railroads, broadly impact whole ecosystems. Roads and railroads can have marked effects on ecological communities. Vehicle collisions are responsible for substantial mortality in animals (Trombulak and Frissell, 2000; Fahrig and Rytwinski, 2009; Becker and Grauvogel, 1991; Gundersen and Andreassen, 1998; Popp et al., 2018), and vehicle traffic elicits avoidance responses in a wide variety of taxa (Prokopenko et al., 2016; Scrafford et al., 2018). Roads can also serve as barriers to movement and disrupt population connectivity (Strasburg, 2006; Shepard et al., 2008). The repeated clearing of roadway margins also creates edge habitat and maintains large areas of habitat in earlier successional stages than surrounding habitats, which can attract species with matching habitat requirements (e.g., for open grass- or shrubland; Forman and Alexander, 1998; Meunier et al., 2000). Such roadside habitat changes, as well as road noise and general disturbance by moving vehicles, has altered the distribution, abundance, and behavior of many species (Meunier et al., 2000; Fahrig and Rytwinski, 2009; McClure et al., 2013). Carrion is also often disproportionately abundant along roads and railways due to vehicle collisions (Becker and Grauvogel, 1991; Gundersen and Andreassen, 1998; Trombulak and Frissell, 2000; Fahrig and Rytwinski, 2009; Popp et al., 2018), which can subsequently attract scavengers and opportunistic predators (Prosser et al., 2008; Lambertucci et al., 2009; Santos and Carvalho, 2011). These conditions generally create a unique species assemblage and an associated set of resources and risks; that risk is often imposed on both predator and prey.

Despite roadway mortality and mortality associated with transmission lines (i.e. collision and electrocution) being substantial sources of anthropogenic mortality in birds (Loss et al., 2015), work to understand the effects of linear features on individual animal movement has been almost entirely restricted to movement of large mammals. Consequently, the effects of linear features on on individual- and population-level avian movement are largely unknown. Their effects on some large birds (e.g., eagles and vultures) are of particular interest for conservation, considering these species are long-lived with slow reproductive rates, and even small amounts of additional anthropogenic mortality may not be sustainable. Many of these species also use carrion as a source of food, a potential attractant to linear features that comes with an increased risk of vehicle collision.

The migration period is already physiologically taxing and associated with elevated mortality in many birds (Newton, 2008; Harrison et al., 2011; Klaassen et al., 2014). While birds differ from landbound taxa in that they should be able to avoid vehicle collisions by flying over roadways and railways, it is unclear whether these features affect the movement or behavior of migrant birds in other ways, such as through habitat modification, changes in prey or carrion distribution and abundance, and/or use of linear features as migration corridors. Given that habitat and potential food resources are important to how a bird uses space during migration (Gill, 2007), there are several biological reasons to expect linear features to alter movement of avian taxa across the landscape.

While effects of habitat and food are important drivers in individual- and population-level movement of migrant birds (Bildstein, 2006; Gill, 2007), the process is more complicated for soaring migrants due to their ability to use air currents to subsidize the energetic costs of flight (i.e. with wind and uplift). The effects of meteorology, especially air currents that develop due to pressure gradients in the atmospheric boundary layer, on movement metrics in many soaring birds are well established (Pennycuick, 1971; Alerstam, 1979; Spaar and Bruderer, 1997; Pennycuick, 1998; Duerr et al., 2012; Lanzone et al., 2012; Duerr et al., 2015; Katzner et al., 2015; Vansteelant et al., 2015; Miller et al., 2016; Shamoun-Baranes et al., 2016; Rus et al., 2017), with mechanistic links between meteorology and movement decisions also demonstrated (Eisaguirre et al., 2018). These atmospheric flight subsides are major components of a soaring bird’s energy landscape (Shepard et al., 2013); however, such subsidies only offset energetic expenses with kinetic energy. Consequently, the distribution of available food energy, which is likely affected by linear features, remains an important component of a soaring bird’s energy landscape, in terms of energy acquisition. Thus, we should expect movement of soaring birds to be dynamically affected by both food resource distributions and available atmospheric flight subsidies (Shepard et al., 2011).

As tools for understanding animal movement, step selection functions (SSFs) have emerged as powerful and robust analytical methods (Fortin et al., 2005; Forester et al., 2009; Potts et al., 2014a,b; Thurfjell et al., 2014; Avgar et al., 2016; Hooten et al., 2017), and when appropriately implemented are able to parse the importance of these different effects on movement decisions. Recent advances allow practical population-level inference while considering individual-level variability in selection and movement (Craiu et al., 2011; Muff et al., 2018). Still, SSFs typically assume that attraction toward or away from different habitats is statistically stationary—that selection does not change through time depending upon an animal’s behavioral or physiological state. Animal behavior and physiology, however, changes across temporal and spatial scales, and attraction or avoidance of different habitats is unquestionably dynamic. Although this is widely recognized (Thurfjell et al., 2014; Hooten et al., 2014; Avgar et al., 2016; Gurarie et al., 2017), behavioral changes are rarely accounted for in any habitat selection analyses, including SSFs, due to the additional complexity models require to capture the non-stationary condition (but see Avgar et al., 2016).

As an animal’s behavioral state changes over time and space, the availability and utility of different habitats across the landscape will also change (Hooten et al., 2014). For example, a migrating animal has an evolved life-history constraint such that it must make migration progress towards a seasonal home range, and, consequently, movements that do not afford progress should be relatively infrequent for that animal. In contrast, while stopped over during migration for foraging or resting, an animal’s movements and habitat selection should change substantially due to differences in energy and habitat requirements during stopover; such a behavioral change would result not only in different use of habitat, but also change in habitat preferences as well. How soaring birds budget behavior and movements, ranging from stopover to migratory, each day across a migration has been shown to be driven by the spatiotemporally-explicit state of the atmosphere (Eisaguirre et al., 2019; Miller et al., 2016), so it is reasonable to suspect that the state of the atmosphere, in part, also influences a soaring migrant’s time-dependent use of the landscape, step selections, and movement decisions. Within such behaviorally-specific use of the landscape, use of linear features could also emerge as being behavioral state- and weather-dependent.

Here, we used a migratory population of golden eagles *Aquila chrysaetos*, a large soaring raptor, as a model system and implemented a novel, biologically justified SSF to investigate how the movement of a long distance migratory soaring bird is affected by both natural and anthropogenic linear features along migration routes. Our SSF incorporated key biologically relevant processes affecting both the energy and resource landscapes, including how soaring migration is driven strongly by wind and uplift conditions. This allowed careful testing of competing hypotheses regarding how terrain and vegetation likely influence movement on individual and population levels. Importantly, this SSF framework allowed us to explicitly show the additional effects linear features can have on space use after accounting for both foraging habitats and dynamic energy landscapes, even despite possible coincidental alignment of migration routes with linear features.

## Methods

### Model system

Golden eagles are a large, long-lived, soaring raptor distributed across the Holarctic (Watson, 2010). Most individuals that summer and breed at high latitudes are long-distance migrants (Watson, 2010; Kochert et al., 2002). Golden eagles are opportunistic predators, capable of using many taxa for food resources, ranging from small mammals and birds to ungulates, and often scavenging carrion (Kochert et al., 2002; Watson, 2010). The population we studied summers primarily in the western Alaska Range and Talkeetna and Chugach Mountains of Alaska, USA and overwinters in the Rocky Mountain West, including Colorado, Utah, Wyoming, Montana, Idaho, Oregon, and Washington, in the US and mid to southern Alberta and British Columbia in Canada (Fig. 1; Eisaguirre et al., 2019; Bedrosian et al., 2018).

**Figure 1:**
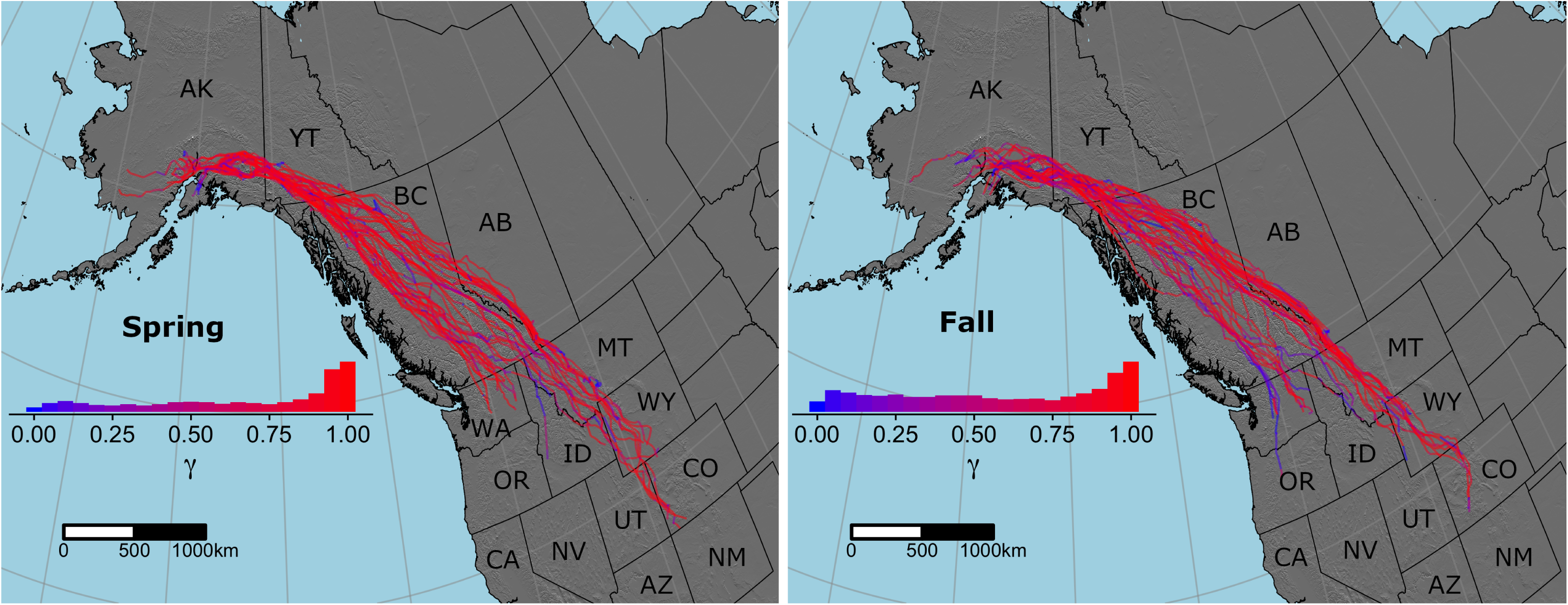
Map of 45 spring and 39 fall individual golden eagle migrations recorded with GPS transmitters in western North America 2014-2017. Insets show frequency distributions of *γ*_*i*_, a time-varying latent variable driven by flight subsidies in a CRW movement model. *γ*_*i*_ close to one (red) indicates more directed, larger-scale migratory movements, and *γ*_*i*_ close to zero (blue) more tortuous, smaller-scale stopover movements.

### Telemetry data collection

We captured golden eagles with a remote-fired net launcher placed over carrion bait near Gunsight Mountain, Alaska (61.67°N 147.35°W). Captures occurred during spring migration, mid-March to mid-April 2014-2016. Adult and sub-adult eagles were equipped with 45-g back pack solar-powered Argos/GPS platform transmitter terminals (PTTs; Microwave Telemetry, Inc., Columbia, MD, USA). Eagles were sexed molecularly and aged by plumage.

PTTs were programmed to record GPS locations on duty cycles, ranging from 8-14 fixes per day during migration, depending on year of deployment. In 2014, PTTs were set to record 13 locations at one-hour intervals centered around solar noon plus a location at midnight local time. 2015 PTTs were programmed to record 8 locations with one-hour intervals centered around solar noon, and in 2016 we revised our programming approach so that PTTs took eight fixes daily with a fixed 3-hr time interval. Poor battery voltage in fall, winter, and spring (September to March) occasionally resulted in PTTs failing to take all programmed fixes, so the resulting GPS tracks had missing observations. Note that such irregular sampling schedules preclude the use of many discrete-time analytical techniques (e.g., conventional and integrated step-selection analyses; Avgar et al., 2016; Hooten et al., 2017).

### Energy landscape step selection function

Step selection functions (SSFs) typically take the form of a separable model, the product of a selection-independent movement kernel and a time invariant selection function:

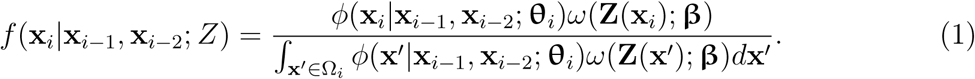

*f* (·) is the marginal probability density of **x**_*i*_, the location of the animal at time *t*_*i*_, given that the animal arrived there after moving from **x**_*i*−2_ to **x**_*i*−1_ over resource field *Z. ϕ*(·) is the selection-independent movement kernel, characterized by movement parameters **θ**_*i*_ and describing how the animal would move over a homogeneous landscape *Z* (Forester et al., 2009). *ω* (·) describes how the animal preferentially selects resources in *Z* based on weights **β** and typically takes a log-linear form (Forester et al., 2009):

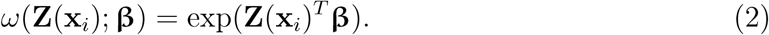

**Z** (**x**_*i*_)^*T*^ is the transpose of a vector-valued function that returns the resource values of interest at **x**_*i*_, and **x**′ is any point in Ω_*i*_, the domain of space available to the animal at *t*_*i*_. *ϕ*(·), along with many modern models for animal movement, is typically a discrete-time correlated random walk (CRW) parameterized in terms of polar coordinates (or step lengths and turn angles; Patterson et al., 2017; Hooten et al., 2017). Such models present challenges, though, in dealing with unequal time intervals between animal locations; step lengths can be normalized by time, but there is not an analogous operation for turn angles. Irregular observations are often handled in the observation equation of discrete-time state-space models; however, with GPS data, we can typically assume negligible observation error and save considerable model complexity by modeling the observations directly with the movement equation (Patterson et al., 2008, 2017; Hooten et al., 2017). Notably, parameterizing a CRW in terms of displacement vectors in a continuous-time framework allows for straightforward relationships with time without an observation equation (Auger-Méthé et al., 2017; Gurarie et al., 2017; Eisaguirre et al., 2019; Jonsen et al., 2019).

To account for behavioral heterogeneity, its predictors, and irregular observations, we implemented our SSF with the following movement model representing *ϕ*(·) (Auger-Méthé et al., 2017; Eisaguirre et al., 2019; Jonsen et al., 2019):

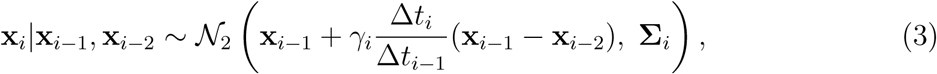

where

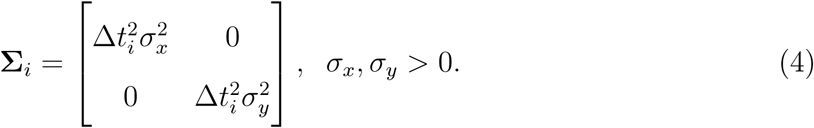

Here, Δ*t*_*i*_ = *t*_*i*_ − *t*_*i*−1_ represents the time interval between Cartesian coordinate vectors **x**_*i*_ and **x**_*i*−1_ for the observed locations of the animal at times *t*_*i*_ and *t*_*i*−1_, and *i* = 1, 2, …, *N* for a track with *N* observations. Note that the movement parameters in equation 1 are **θ**_*i*_ = (*γ*_*i*_, **Σ**_*i*_). The latent variable *γ*_*i*_ correlates displacements (or ‘steps’) and can be interpreted to understand the type of movement, and thus behavior, of migrating individuals: estimates of *γ*_*i*_ closer to one indicate directionally-persistent, larger-scale migratory movement, while estimates of *γ*_*i*_ closer to zero indicate more-tortuous, smaller-scale stopover movement (Breed et al., 2012; Auger-Méthé et al., 2017; Eisaguirre et al., 2019; Jonsen et al., 2019). The behavioral process can be written

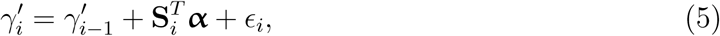

where

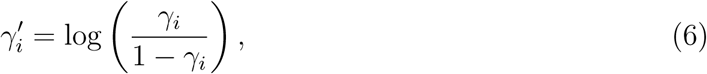

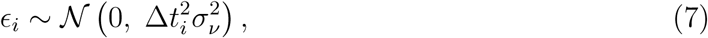

and **S**_*i*_ is the vector of environmental covariates at location **x**_*i*_ and time *t*_*i*_. Each element of ***α*** is an estimated parameter representing the magnitude and direction of the effect of its respective covariate on *γ*_*i*_ in addition to the effect of *γ*_*i*−1_. Including *γ*_*i*−1_ here specifies explicit serial correlation in the behavioral process so that any environmental effect is not overestimated.

### Inference

Practical inference with SSFs often requires estimating the movement process *ϕ*(·) and selection function *ω* (·) separately (Fortin et al., 2005; Forester et al., 2009; Potts et al., 2014b,a; Thurfjell et al., 2014; Hooten et al., 2017). Here, doing such corresponds to first making inference about the animal’s movement and behavioral processes in addition to effects of environmental covariates on those processes (equations 3-7). We then proceed to estimate effects of habitat features that could additionally affect space use through the animal’s preferential selection (equation 2). Although, estimating *ϕ*(·) and *ω* (·) independently could affect inference of respective parameters, it has been shown to have little to no effect on *ϕ*(·) (Potts et al., 2014b), and there are ways to minimize bias in *ω* (·) (*sensu* Forester et al., 2009).

We fit our movement model (equations 3-7), representing *ϕ*(·) in equation 1, in a Bayesian framework with Stan in R (Stan Development Team, 2018; R Core Team, 2018), following Eisaguirre et al. (2019), with five chains of 200,000 Hamiltonian Monte Carlo (HMC) iterations, including 100,000 for warm-up, and retaining 10,000 samples for inference (see Eisaguirre et al. (2019) and/or the code provided as supporting information for prior choice). Fitting *ϕ*(·) was done independently of *ω*(·) for each individual migration.

The integral in the denominator of equation 1 is essentially always computationally prohibitive. However, a number of approximate methods have been proposed (see Hooten et al. 2017). To estimate an SSF with an use-availability design, *k* available steps with endpoints 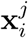 for *j* = 1, 2, …, *k* matched to the move from **x**_*i*−1_ to **x**_*i*_ are generated from *ϕ*(**x**|**x**_*i*−1_, **x**_*i*−2_; **θ**_*i*_). Then, the resource vectors **Z**(**x**_*i*_) and 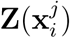 are populated from the appropriate data sources. We chose *k* = 5 for our analysis (Thurfjell et al., 2014), and under the Bayesian paradigm, simulating from our fitted *ϕ*(**x**|**x**_*i*−1_, **x**_*i*−2_; **θ**_*i*_) is analogous to sampling from the conditional posterior predictive distribution (Hooten et al., 2014, 2017)— the probability of a new *i*th observation given the observed data—which, for a ‘new observation’ 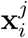, we deno 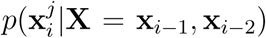, where **X** is the *N* × 2 matrix containing each **x**_*i*_ (Fig. 2). Sampling from each 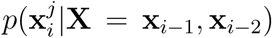 has the advantage of accounting for all parameter uncertainty and is fairly simple in the most commonly used Bayesian modeling languages (e.g., Stan and BUGS). Note that 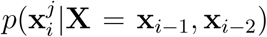 is (analytically) the result of integrating over the model parameters, including the time-varying latent behavioral variable *γ*_*i*_, so each 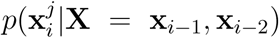 is conditioned on the animal’s behavior at time *t*_*i*_. Finally, to estimate **β**, the comparison of used and available steps for each animal is carried out with conditional logistic regression. Hooten et al. (2014) present a similar approach that leverages the posterior predictive of a continuous time CRW fit with a Kalman filter (Johnson et al., 2008) to estimate **β** based on the smoother (use) and predictor (available) distributions. An advantage to their method is handling observation error; however, since observation error is negligible in our case, we decided to characterize use with the observed data, rather than a predicted distribution.

**Figure 2:**
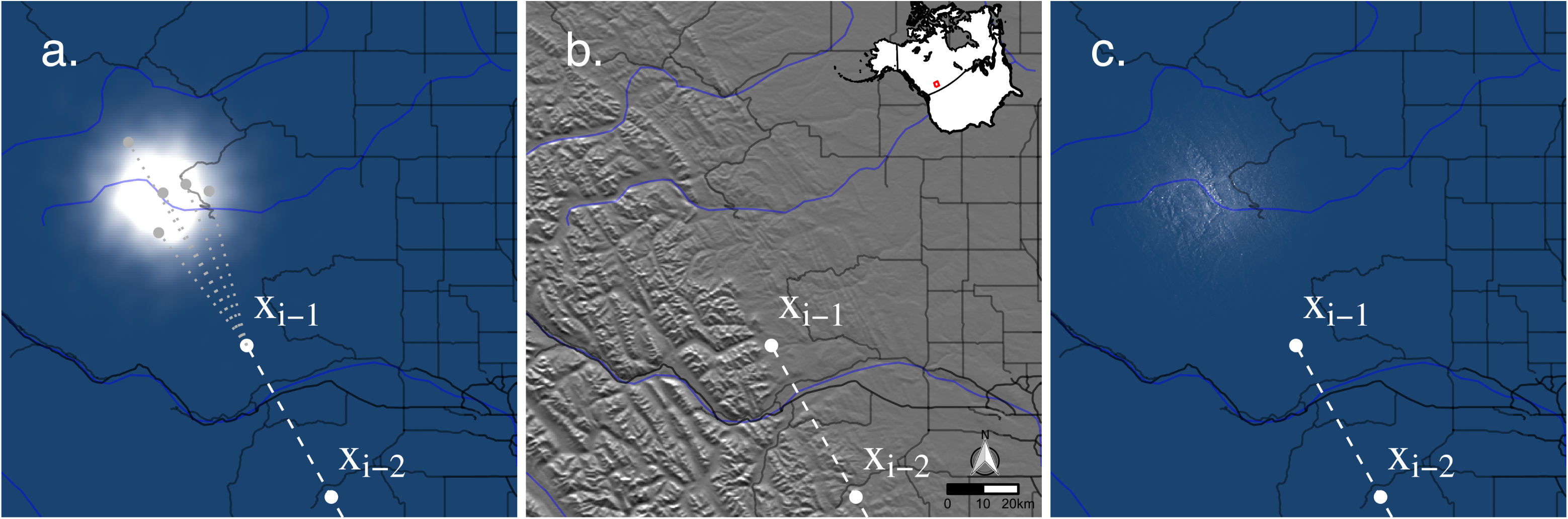
Example movement kernels from an el-SSF fitted to golden eagle migrations 2014-2017 conditioned on the displacement vector from **x**_*i*−2_ to **x**_*i*−1_. (a) 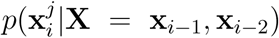 is the conditional posterior predictive distribution of the fitted selection independent kernel *ϕ*(**x**_*i*_|**x**_*i*−1_, **x**_*i*−2_), (b) is an elevation map showing roads and waterways, representing aspects of the landscape *Z*, and (c) 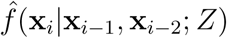 is the product of the fitted selection independent kernel and fitted habitat weighting function. Grey points show example available points 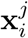 drawn from 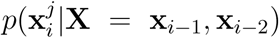 for use in estimation of *ω* (·) with the use-availability design. Note that both (a) and (c) show spatiotemporally explicit probability densities for **x**_*i*_, a golden eagle’s location during migration at time *t*_*i*_ given that the eagle traveled from **x**_*i*−2_ to **x**_*i*−1_ with (estimated) correlation *γ*_*i*_ ≈ 1 driven by uplift.

In presenting our energy landscape SSF (el-SSF), above, we did not account for multiple individuals, but this was just for notational simplicity. We estimated habitat selection parameters hierarchically across individuals. Hierarchical conditional logistic regression to estimate individual- and population-level effects presents estimation challenges (Duchesne et al., 2010), so we chose to implement a Poisson approximation of the hierarchical case, which allows Bayesian inference with integrated nested Laplace approximations (INLA; Muff et al., 2018). We followed Muff et al. (2018) and used R and the package `r-INLA` to estimate the resource weights **β** in *ω* (·) (Rue et al., 2009; R Core Team, 2018). See Muff et al. (2018) for prior choice. A 3-knot linear spline with knot locations at the quartiles of the step lengths was included in each candidate selection model to minimize bias in estimating **β**, as 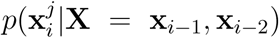 is not actually selection-independent (Forester et al., 2009). Parameterizing a movement kernel with real data that is truly selection-independent is essentially impossible, given that we cannot observe animals moving over a homogeneous *Z* landscape (but see Avgar et al., 2016).

As golden eagles are diurnal, we chose to only include daytime movements, which we defined as those between sunrise and sunset, in estimating the el-SSF. Sunrise and sunset times local to each GPS point were calculated with the R package `maptools` (Bivand and Lewin-Koh, 2016).

### Candidate models of movement & selection

We proposed a set of candidate models of movement and habitat selection and compared them using the widely applicable information criterion (WAIC; Watanabe, 2010), which is calculated by `r-INLA`, asymptotically equivalent to Bayesian cross validation, and an improvement over the deviance information criterion (Gelman et al., 2014). Each model represented a hypothesis for how golden eagles move and select for space during migration. Both the environment and the internal state of eagles varies substantially between spring and fall, so we fit the el-SSF and ranked candidate models independently for each season. A number of the variables included in our models were temporally dynamic, so much of any inter-annual variation was captured implicitly.

Given that golden eagle flight and behavior is driven by atmospheric flight subsidies (Duerr et al., 2012, 2015; Katzner et al., 2015; Miller et al., 2016; Rus et al., 2017; Eisaguirre et al., 2019), thermal uplift, orographic uplift, and wind support were included in all candidate models (in *ϕ*(·); see below) to account for the dynamic energy landscape. Before comparing models including effects of anthropogenic linear features, we first wanted to determine which natural variables are most important to habitat selection, so we constructed a set of six candidate models, in addition to a ‘no selection’ null model, that generally corresponded to effects of the following: terrain, landcover, terrain + land-cover, terrain + waterways, landcover + waterways, and terrain + landcover + waterways. Comparing these models first allowed us to pare down the set of biologically plausible models that might otherwise be quite large if all habitat and linear feature variables were considered together.

We characterized terrain with elevation and slope aspect. We suspected that eagles would select for higher elevations, and given that conditions at higher elevations vary strongly along a latitudinal gradient, elevation was included in models as an interaction with latitude. South-facing slopes are exposed to more intense solar radiation, and thus produce more thermal uplift. Although thermal uplift is a favored energetic subsidy (Duerr et al., 2012), changes in urgency, especially as individuals approach the breeding grounds, might lead them to forego use of south-facing slopes in favor of a more direct route (Miller et al., 2016). Additionally, ambient conditions (e.g., prevailing air temperature/pressure) generally change substantially with latitude, which could affect the degree to which eagles favor south-facing slopes as a source of uplift. We thus also included aspect as an interaction with latitude. We considered including terrain ruggedness; however, it was highly colinear with elevation, so it was not included.

Landcover was characterized by vegetation and snow cover. Prey availability likely varies with vegetation and snow, but densely vegetated areas could generate thermal up-lift (Howard and Stull, 2013). In the field while capturing eagles during spring migration, we observed eagles seemingly thermal soaring over areas predominantly flat and densely covered with dark vegetation (i.e. *Picea* spp.) comprising an otherwise snow covered landscape, which would typically not offer thermal uplift.

Waterways were treated as natural linear features. Waterways would be a source of prey (i.e. waterfowl), but could also be used for navigation. Golden eagles that migrate to Alaska have been shown to use the long ‘trenches’ (very straight, long valleys) in the Canadian Rocky Mountains (Kochert et al., 2002; McIntyre et al., 2008; Eisaguirre et al., 2019). Large bodies of water could also be barriers to movement (Kochert et al., 2002).

After selecting the best approximating model given our set of models that included natural covariates, we generated another set of three candidate models that included anthropogenic linear features; the two types considered were roads and railroads. Given that collisions with vehicles and transmission lines along roads are a leading cause of anthropogenic avian mortality (Loss et al., 2015), we suspected eagles might avoid them due to mortality risk. Alternatively, roads likely provide carrion, which could attract eagles, perhaps even to follow them during migration. Railways have not been documented as a major source of mortality risk to raptors and other birds—although a juvenile eagle banded in Alaska was killed by a train (T. Booms, *unpubl. data*). Railways are, however, responsible for substantial mortality in large mammals (e.g., ungulates; Becker and Grauvogel, 1991; Gundersen and Andreassen, 1998; Popp et al., 2018), potentially concentrating carrion resources for migrating eagles. As both railway and road densities decline with increasing latitude in North America, they were included in models as an interaction with latitude in addition to their main effects.

We were interested in investigating effects of transmission lines on eagle movement as they are a leading cause of raptor mortality (Loss et al., 2015). However, power line corridor data are largely proprietary and confidential and thus were not available to include in models. We were also interested in the effects of wind energy developments (Pagel et al., 2013; Loss et al., 2015); however, we did not include them in candidate models due to their apparent minimal availability to the eagle migrations we sampled. We present summary statistics regarding interactions between eagles and wind energy developments in Appendix S1.

### Covariate data

#### Flight subsidies

We gathered meteorological flight subsidy data for eagle tracks with the Track Annotation Service in Movebank (Dodge et al., 2013). These variables were introduced as covariates **S**_*i*_ into the behavioral process in the the movement kernel *ϕ*(·) (Eisaguirre et al., 2019). In the el-SSF, flight subsidies **S**_*i*_ coupled with **α** modify what is available to an eagle at each *t*_*i*_ by driving the behavioral dynamics (equations 5-7).

Thermal uplift was bilinearly interpolated from European Centre for Medium-Range Weather Forecasts (ECMWF) reanalyses, and orographic uplift from the nearest neighbor (grid cell) by pairing National Center for Environmental Predictions (NCEP) North American Regional Reanalysis (NARR) data with the Advanced Spaceborne Thermal Emission Reflection Radiometer (ASTER) Global Digital Elevation Model (GDEM; Brandes and Ombalski, 2004; Bohrer et al., 2012). Wind data were interpolated bilinearly from the NCEP NARR *u* (westerly/zonal) and *v* (southerly/meridional) components of wind, from which we calculated the wind support (Safi et al., 2013).

#### Terrain & habitat

Elevation, slope, vegetation, and snow cover data were gathered for all locations **x**_*i*_ and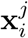 with the Env-DATA system (Dodge et al., 2013). The source of the elevation and slope data was the ASTER. Env-DATA provides the *u* and *v* components of slope. We used the *v* (south to north) component as it represents the degree to which the slopes in the grid cell for each **x***i* and 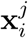 are south-facing (hereafter slope southing). Percent vegetation and snow cover of grid cells were predicted with the NCEP NARR.

#### Linear features

Waterway data were gathered from the Commission for Environmental Cooperation 2009 Lakes and Rivers dataset. Road and railroad data were gathered from the United States Geological Survey National Transportation dataset and Canadian National Road/Railway Network. The road types included in models were those considered arterials, (state or county) highways, and freeways.

These data were included in candidate models of selection as distance to nearest (waterway, road, or railroad) measured at step endpoints. All distances were measured in R with the package `sp` (Pebesma and Bivand, 2005; Bivand et al., 2013).

We chose to include only endpoint effects in candidate models of selection for two main reasons. First, irregular observations lead to inconsistent uncertainty between observed locations. Interpolations between observed points are less accurate with larger Δ*t*_*i*_, so normalizing by Δ*t*_*i*_, although simple, would not be entirely appropriate; other assumptions would be required (Thurfjell et al., 2014). Second, migrants are essentially required (by their life history) to cross linear features that are not precisely parallel to movement during migration, so movements that do not cross roads, for example, would be minimally available, making quantifying any effect of crossings on selection difficult.

Lastly, we conducted a brief simulation study, which we present in Appendix S3, to validate that the el-SSFs detected real effects of linear features. This was to ensure that apparent selection for linear features as estimated within the SSF framework was not simply an artifact of the distribution of linear features on the landscape, even if they coincidentally aligned with migration routes.

## Results

Of the tags deployed, 32 provided at least one migration with sufficient data to estimate the movement kernel *ϕ*(·). Nine were deployed on females and 23 on males, and all were adults (at least five years old). From those individuals, 17,386 realized (used) daytime steps were included in the step selection analysis spanning 45 spring and 39 fall individual migrations 2014-2017. Median (interquartile range) spring and fall departure dates across years were 10 March (6 days) and 2 October (11.5 days), respectively, and arrival dates were 30 March (6 days) and 13 November (20 days). Only 25 of the 17,386 steps analyzed intersected 1 km buffers surrounding wind turbines; twenty endpoints of the steps were within 5 km of a wind turbine; and only two were within 2 km (Appendix S1).

### Atmospheric flight subsidies

We found thermal uplift to be the main driver of migratory behavior during both spring and fall migrations across years, though there was a high degree of variability in how flight subsidies (wind and uplift) drove behavior among individuals (Fig. 3). Our sample contained more than twice as many males as females, but we found little evidence for an effect of sex on the behavioral process (Fig. 3). While an eagle’s response to flight subsidies did not change markedly between seasons (Fig. 3), eagles tended to adjust their pace to move more quickly in the spring by budgeting more time to faster, directed movement. During fall migrations, eagles moved more slowly, and budgeted comparatively more time to slower, less directed movement resembling searching and stopover (Fig. 1).

**Figure 3:**
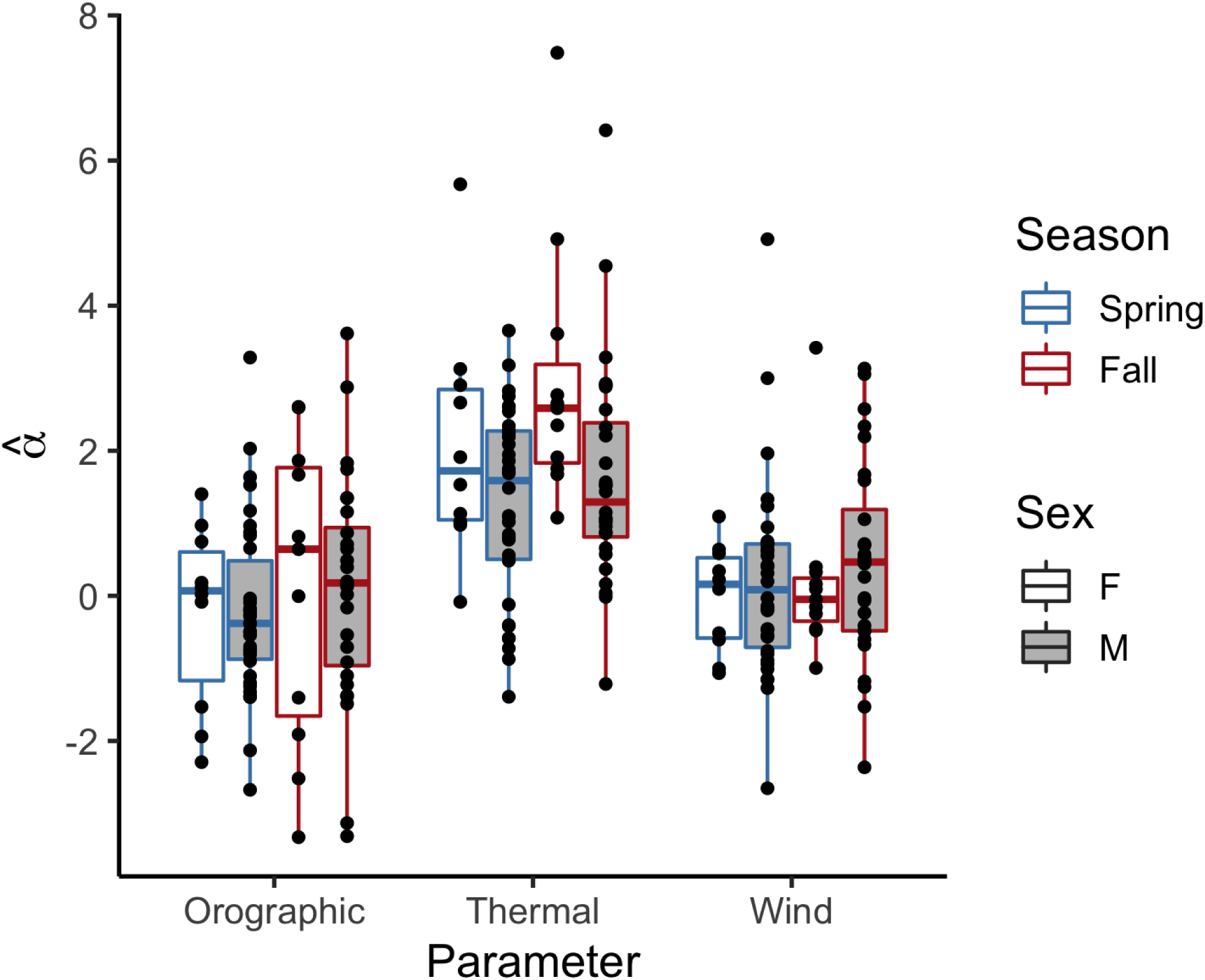
Figure 3: Effects of flight subsidies (orographic uplift, thermal uplift, and wind support) on golden eagle movement during migration 2014-2017 estimated with a CRW movement model. 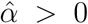 indicates the flight subsidy is associated with more directed, larger-scale migratory movements, and 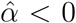 more tortuous, smaller-scale stopover movements.

### Model selection

All habitat variables included in the el-SSF improved fit, though we note that terrain was especially informative, strong evidence that habitat is an important component of the movement process (Table 1). Although the CRW and flight subsidies *ϕ*(·) can predict a relatively large area of high probability for step selection (Fig. 2a), when habitat is considered, the high probability region becomes quite concentrated (Fig. 2c). A model including elevation, slope southing, snow and vegetation cover, and waterways was top ranking in both spring and fall (Table 1). The addition of linear features further improved fit (Table 1), with substantial support for both roads and railroads driving movement in spring and fall in addition to terrain, landcover, and waterways.

**Table 1:**
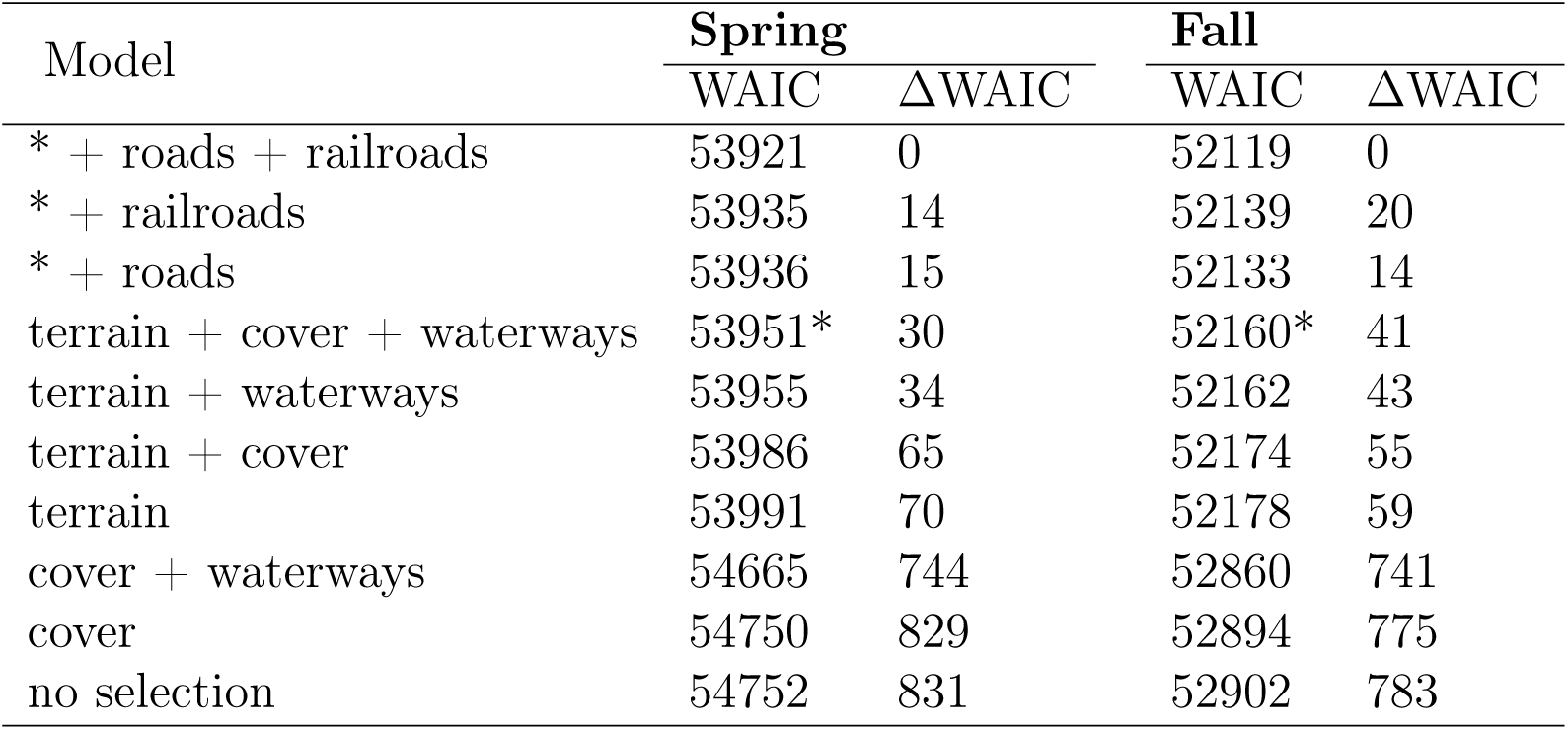
Ranking of candidate models of habitat selection by golden eagles during migration 2014-2017 by the widely-applicable information criterion (WAIC). ‘terrain’ includes elevation and slope southing; ‘cover’ includes percent vegetation and snow cover; and linear features (roads, railroads, and waterways) were included as distance to nearest. * corresponds to the model with only natural variables chosen to include with anthropogenic linear features.

### Natural landscape features

Eagles generally used and preferred lower elevations in the spring and higher in the fall (Fig. 4 & 5), though there was considerable variation among individuals (Fig. 5). In spring, selection for higher elevation decayed with latitude, while in fall, change in elevation preference with latitude was variable across individuals (Fig. 5). All eagles favored south-facing slopes in both seasons, showing a stronger preference with increasing latitude in spring (Fig. 5). Although snow-covered and vegetated areas were used to a greater degree in spring (Fig. 4), all individuals showed a preference for snow-covered and less vegetated areas in spring (Fig. 5).

**Figure 4:**
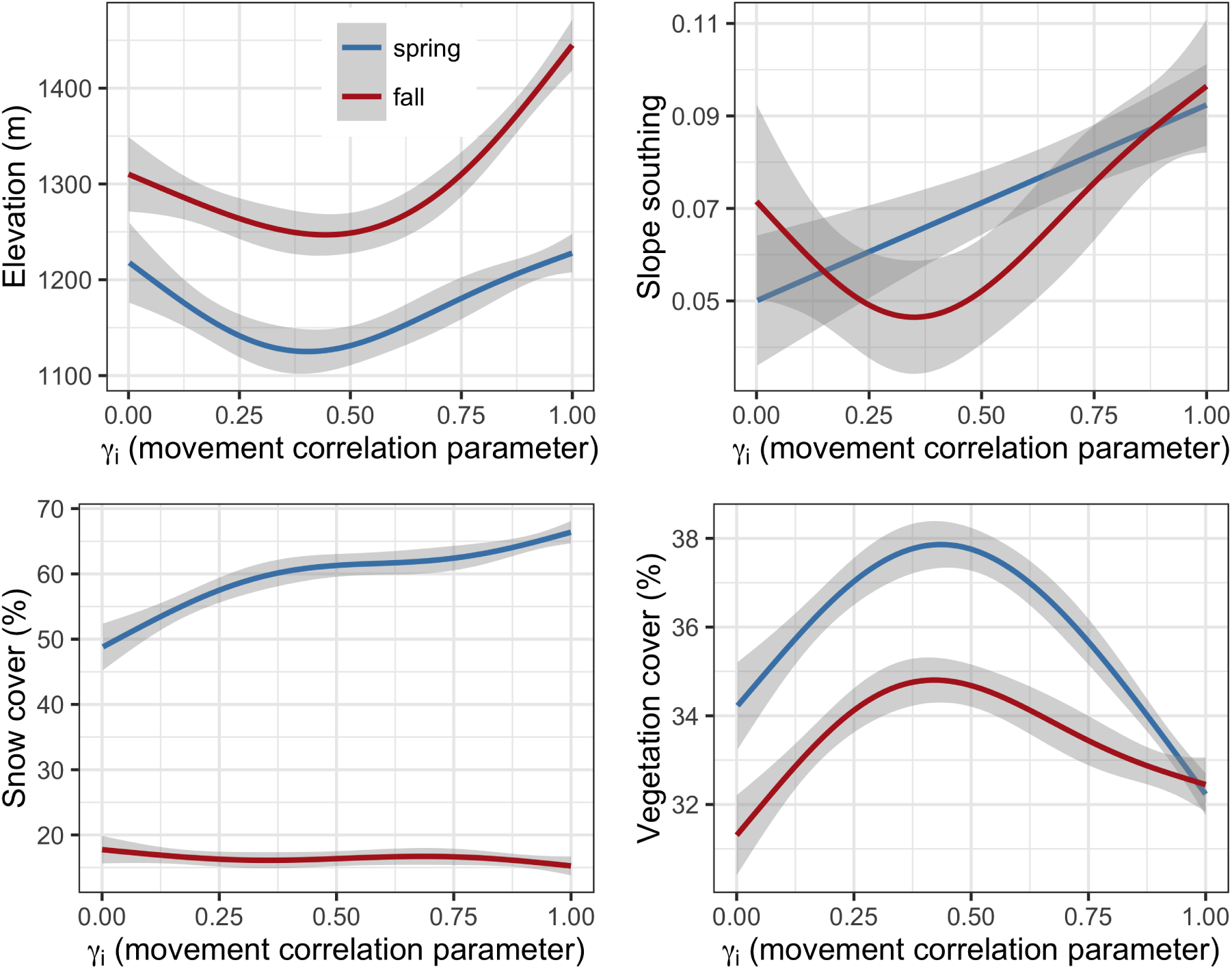
Empirical use distributions smoothed with a generalized additive model (*df* = 4; shaded areas are 95% confidence intervals) over all golden eagle spring and fall migrations 2014-2017 as a function of the movement parameter *γ*_*i*_. *γ*_*i*_ is a time-varying latent variable driven by flight subsidies in a CRW movement model. *γ*_*i*_ close to one indicates more directed, larger-scale migratory movements, and *γ*_*i*_ close to zero more tortuous, smaller-scale stopover movements.

**Figure 5:**
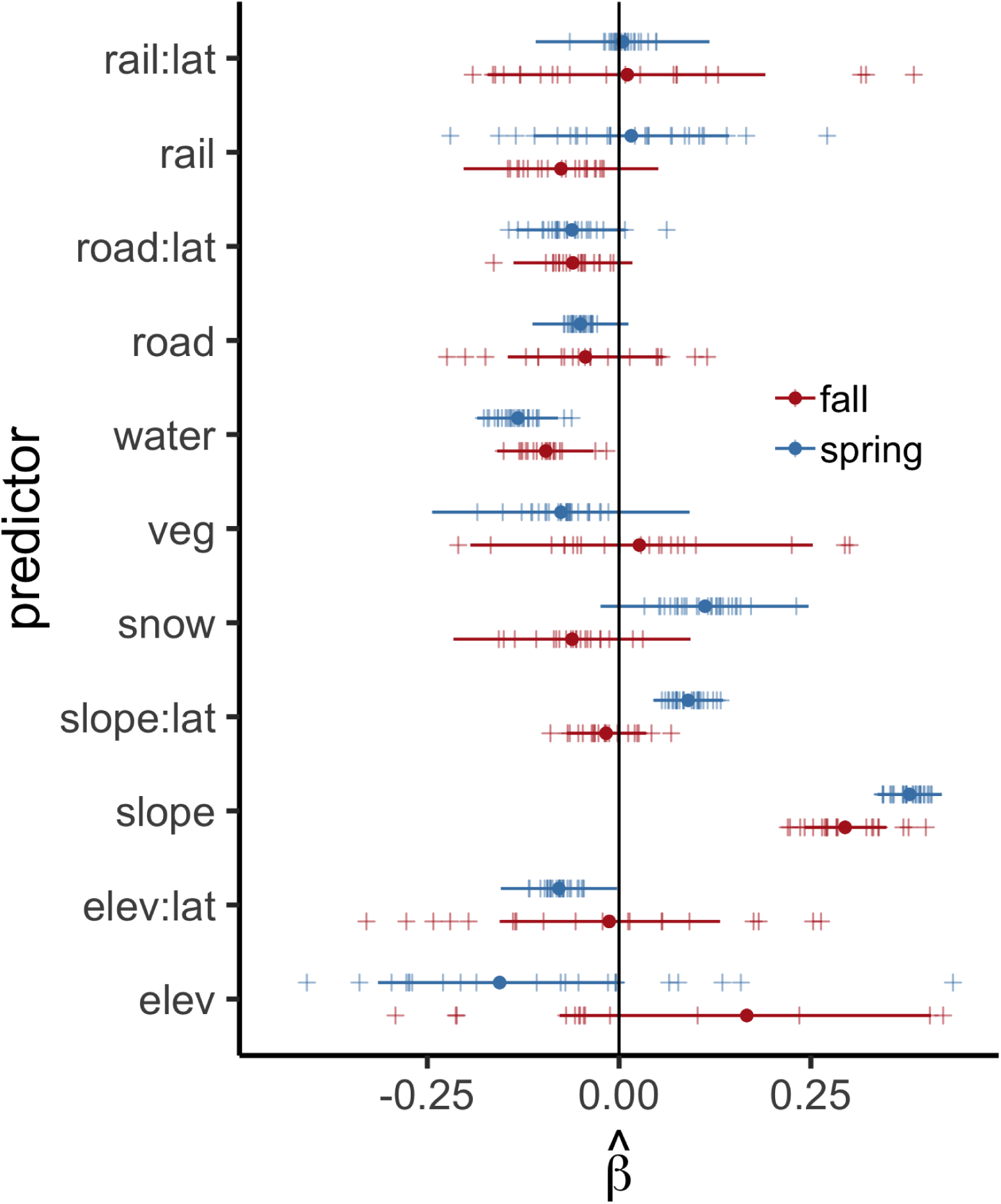
Effects of habitat variables on golden eagle habitat selection during spring and fall migration 2014-2017 estimated with el-SSFs. Points are posterior means and horizontal lines 95% highest posterior density (HPD) intervals for the population-level effects. Crosses correspond to the individual-level posterior means. Predictors included were elevation (‘elev’), slope southing (‘slope’), percent snow cover (‘snow’), percent vegetation cover (‘veg’), and distance to nearest railroad (‘rail’), road (‘road’), waterway (‘water’), some as an interaction with latitude (‘lat’). Negative estimates on distance to nearest linear feature correspond to selection for areas close to the linear feature. Estimates are on a standardized scale. Note that we have shortened the *x* axis here for clarity, but the full version is provided in Appendix S2.

**Figure 6:**
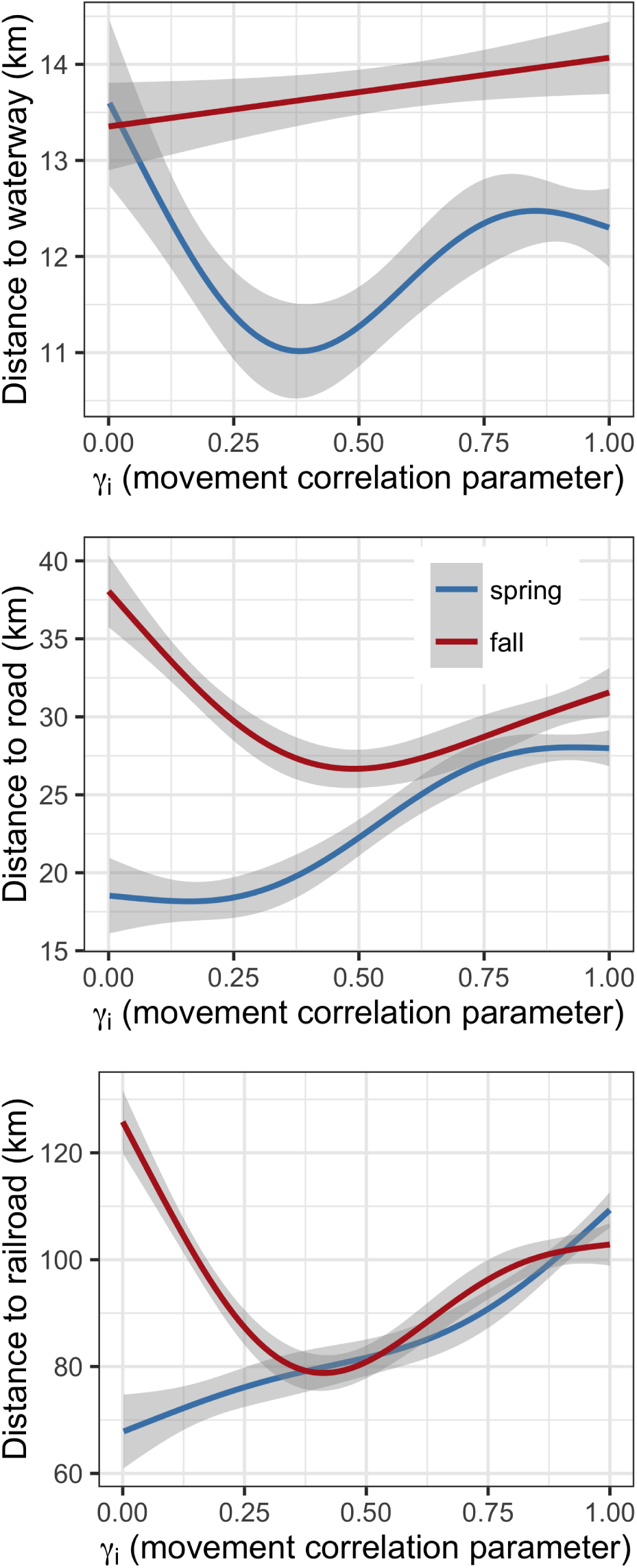
Empirical use distributions for linear features smoothed with a generalized additive model (*df* = 4; shaded areas are 95% confidence intervals) over all golden eagle spring and fall migrations 2014-2017 as a function of the estimated movement parameter *γ*_*i*_. *γ*_*i*_ is a time-varying latent variable driven by flight subsidies in a CRW movement model. *γ*_*i*_ close to one indicates more directed, larger-scale migratory movements, and *γ*_*i*_ close to zero more tortuous, smaller-scale stopover movements.

In both seasons, eagles showed a preference for areas close to waterways given the habitat available (Fig 5–7). However, there was less selection for areas near waterways in the fall compared to spring.

### Anthropogenic linear features

Eagles were often near linear features but more frequently closer to roads and railroads during spring migration than fall (Fig. 8). el-SSF predictions indicated that probability of using a given area increased if that area was closer to a road, especially in spring (Fig 7). Eagles’ preference for areas near roads decayed slightly with increasing latitude (Fig. 5 & 9), and there was considerably individual variability in preference for areas near roads during fall migration (Fig. 5 & 9). Eagles exhibited a slight preference for railroads in fall (Fig. 5 & 9); however, there was, again, substantial variability among individuals in both seasons (Fig. 5 & 9). See Appendix S3 for results of simulations showing SSFs parse actual selection from incidental use of linear features when linear features and movement routes are coincidentally parallel.

**Figure 7:**
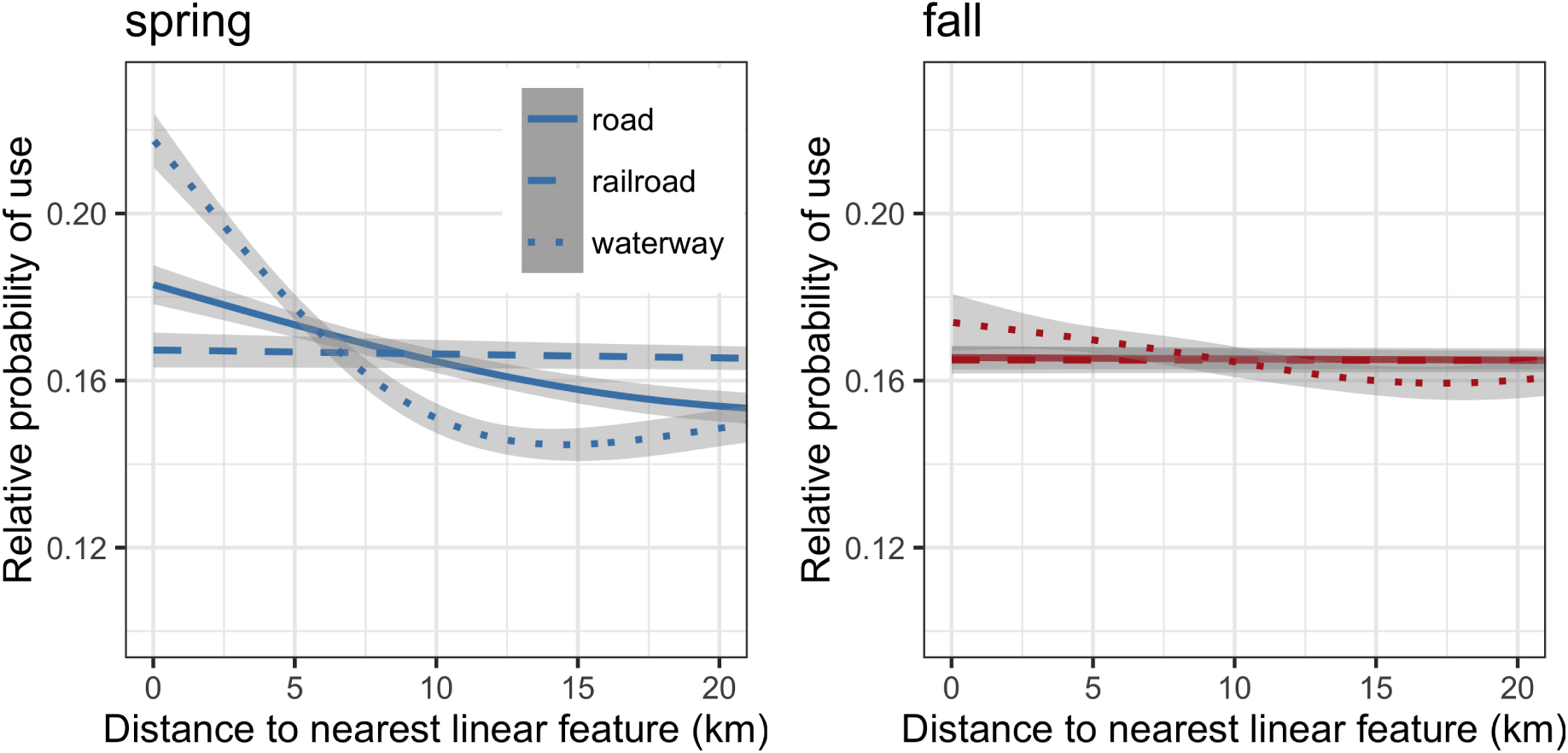
Average effect of distance to nearest linear feature on space use of golden eagles during spring and fall migrations 2014-2017 estimated with el-SSFs. This is conditioned on how habitat was distributed within the availability distribution for the population sampled (Avgar et al., 2017). Curves depict the smoothed (generalized additive model, *df* = 6) nonparametric function between the distance to linear feature and relative probability of use, and shaded areas correspond to 95% confidence intervals.

**Figure 8:**
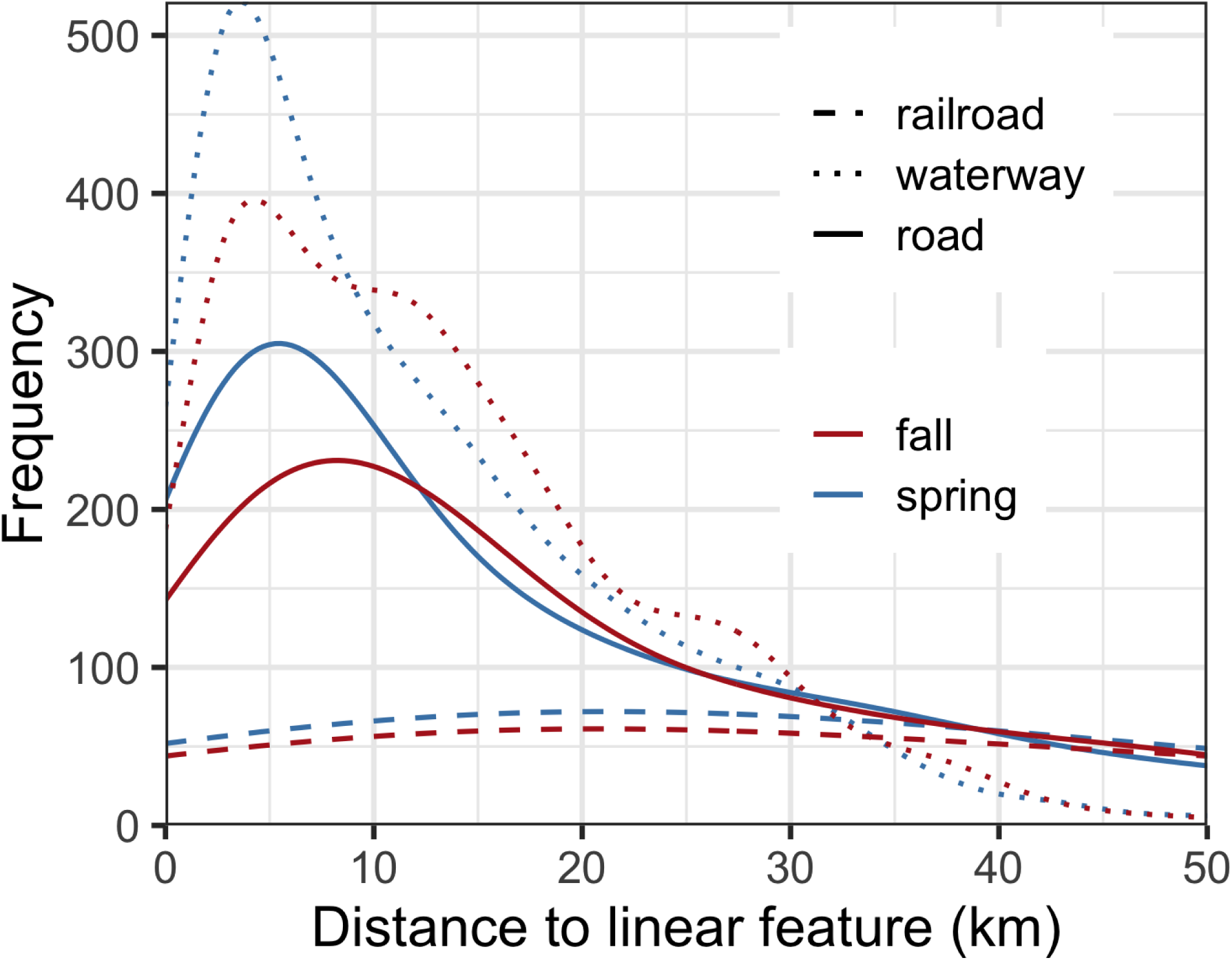
Smoothed empirical distributions of distance from 17,386 daytime golden eagle GPS transmitter locations to the nearest road, railroad, and waterway during spring and fall migrations 2014-2017 in western North America.

**Figure 9:**
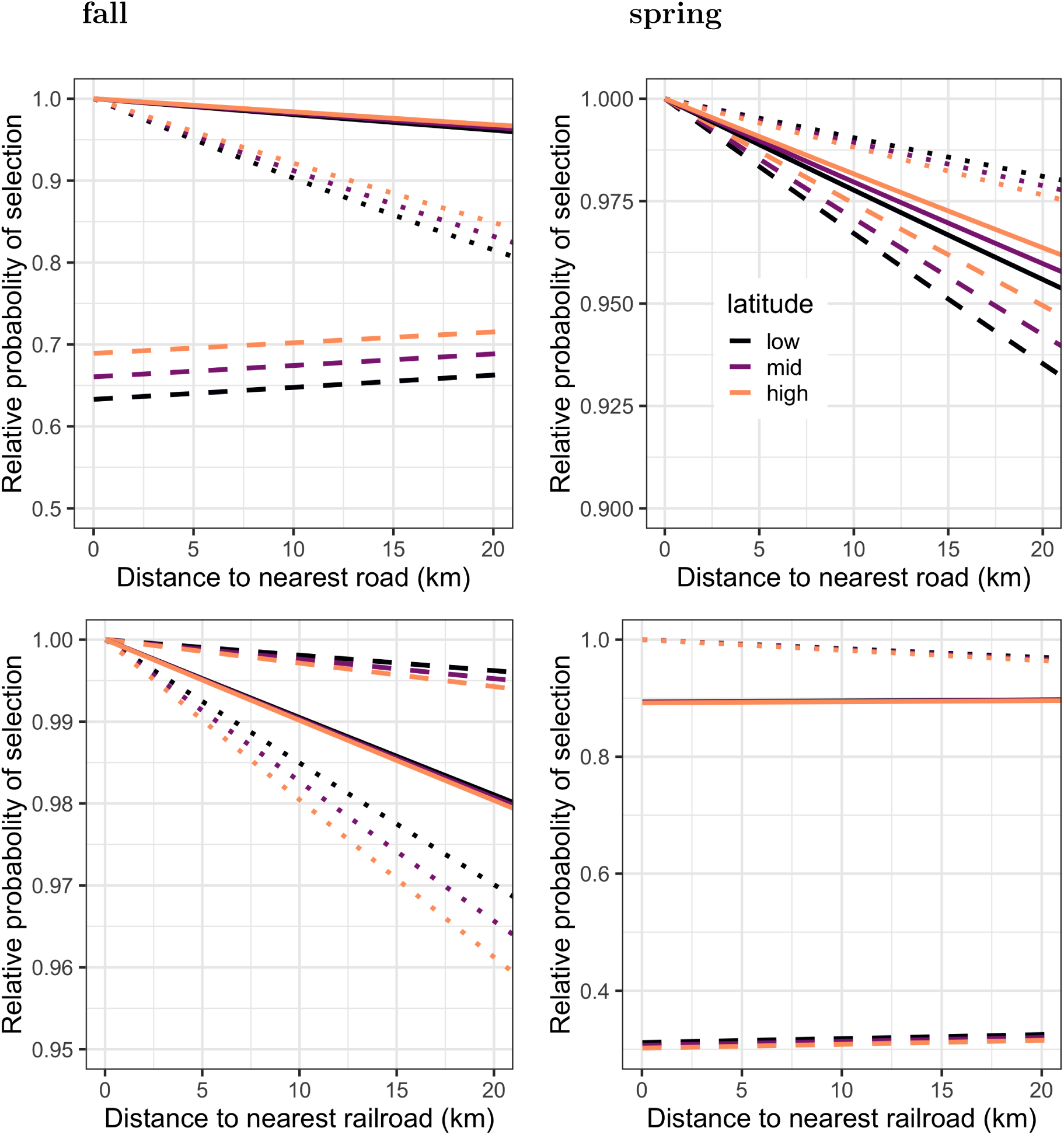
Population-level (solid) and two individual-level (dashed and dotted) preference curves for anthropogenic linear features predicted by an el-SSF fitted to golden eagle migrations 2014-2017. These describe the relative probability of selection by golden eagles during spring and fall migration, assuming road distances are uniformly distributed and equally available to eagles (Avgar et al., 2017). Individual-level curves shown to highlight the variation among individuals. ‘low’, ‘mid’, and ‘high’ latitude correspond to approximately 41°, 51°, and 61°N latitude, respectively.

### Effects of behavior on habitat use

The use of anthropogenic linear features and habitat changed substantially depending upon expressed behavior, as captured by estimates of *γ*_*i*_ (Fig. 4 & 6). When eagles engaged in less directed, more tortuous movements, typically associated with stopover, these behaviors were performed in areas closer to roads and railroads in spring and farther from roads and railroads in fall (Fig. 6). Areas with higher percent vegetation cover were used when eagles were moving with intermediate directional persistence and movement rate, while very low persistence and very high persistence states were associated with less vegetated areas (Fig. 4). Use of space near waterways also varied with behavioral state, such that eagles were closer to waterways while moving with intermediate directional persistence and rate (Fig. 6). During fall migration, eagles generally used higher elevation terrain while making faster-paced, directionally-persistent movements, and during both spring and fall migration, eagles used more south facing slopes while making faster-paced, directionally-persistent movements (Fig. 4).

## Discussion

The effects of linear features on the movement of large terrestrial mammals are relatively well understood (Dyer et al., 2002; Whittington et al., 2004; Dickson et al., 2005; Latham et al., 2011; Whittington et al., 2011; McKenzie et al., 2012; Tremblay and Clair, 2009; DeMars et al., 2016; Dickie et al., 2017; Popp et al., 2018; Dickie et al., 2019), but, here, we showed that anthropogenic linear features can also affect movement and space use of avian species during migration. Our approach was analytically nuanced, incorporating the affects of energy subsidies key to soaring movement into the the movement process. This helped disentangle the effects of such subsidies from habitat selection, which would not have otherwise been possible with more phenomenological models, such as conventional resource or step selection functions. Specifically, the el-SSF accounted for favorable up-lift and wind conditions (i.e. a soaring bird’s energy landscape), which probabilistically restricted the habitats available for selection; the effects of linear features could then be additionally estimated to further explain movement and space use (Fig. 2). Such an approach is superior, as it mechanistically restricts where a soaring bird is likely to move based on the energy landscape and weather conditions. A migrant can be less likely to make moves that accrue several hundred kilometers of migration progress during a day when there is limited uplift, for example, so dynamically restricting the el-SSF movement process based on these conditions is a natural extension of the current static, discrete-time CRWs employed in conventional SSFs (e.g., Fortin et al., 2005; Forester et al., 2009; Thurfjell et al., 2014; Potts et al., 2014b,a).

Building movement kernels with more biological relevance—beyond basic random walks—into SSFs has been suggested (Thurfjell et al., 2014; Jonsen et al., 2019), but to our knowledge, this is the first time it has been executed while maintaining the practicality of (hierarchical) conditional logistic regression for estimation of resource selection parameters. The el-SSF also allowed us to account for and detect variation among individuals, which has been shown to be key to correct inference of resource selection patterns (Lesmerises and St-Laurent, 2017), while still making population-level inference of habitat and resource selection. Lastly, we were able to show that use of habitat varied with the expressed behavior (tracked with the dynamic value of *γ*_*i*_, the movement correlation parameter), ranging from slower-paced, stopover movements to faster-paced, migration movements.

These behavioral changes were built into the CRW movement kernel in the el-SSF, allowing us to leverage aspects of related mathematical and movement ecology theory. In spring, eagles used areas closer to roads and railroads while performing slower-paced movements suggestive of migratory stopover (Fig. 6). As movements that tend to keep an animal in the same area (i.e. slower-paced/searching movements) coincide with greater residence time (Turchin, 1991, 1998), we can infer that eagles’ residence time nearer roads and railroads during spring migration is higher compared to other areas on the landscape. While this also implies eagles would be relatively more abundant there at any given time (Odendaal et al., 1989), eagle residence time would be conditional on atmospheric conditions, as stopover movements are more likely when weather conditions do not support thermal uplift (Fig. 3; Eisaguirre et al., 2019; Miller et al., 2016). So, eagles’ use of areas nearer roads and railroads might ultimately vary with uplift conditions. However, as uplift conditions are driven by the daily development of the atmospheric boundary layer—uplift is least available in morning and evening due to limited insolation during those times of day—eagle residence time near linear features may generally be greatest in morning and evening.

### Extending SSFs

The el-SSF approach offers novel analytical utility, but using movement data to investigate some important questions regarding details of animals’ use of linear features requires a statistical method that can assess state-specific selection for different landscape features. An important question, here, is how an eagle might balance scavenging road-killed carrion with the risk of vehicle collision across different behavioral or physiological states. State-specific selection in SSFs is an active area of analytical development (Hooten et al., 2014; Avgar et al., 2016; Hooten et al., 2017; Karelus et al., 2019; Scharf et al., 2019), and while the el-SSF did not explicitly incorporate behaviorally-specific selection into its framework, we were still able to infer behavior-specific use of linear features and habitat plus seasonal variation in those patterns (Figs. 4-6). While progress is being made (e.g., Avgar et al., 2016), these findings further support the need to consider behavioral heterogeneity in resource selection analyses and work towards overcoming related analytical obstacles.

Estimating behaviorally-specific selection coefficients requires introducing movement parameters into both the movement and selection processes, and thus imparts dependence between *ϕ*(·) and *ω*(·) (equation 1). This structure often precludes the use of conditional logistic regression for obtaining unbiased estimates of **β** without special consideration of that dependence (Forester et al., 2009; Avgar et al., 2016).

### Eagle migration & anthropogenic linear features

#### Behaviorally-dependent use of linear features

There are a number of possible reasons for why migrating eagles might select and use areas close to anthropogenic linear features. They may use linear features for (1) navigation, (2) increased movement rate—these features may produce favorable flight conditions—and/or (3) access to food resources. Although other taxa have been shown to use linear features to increase movement rate (Latham et al., 2011; McKenzie et al., 2012; Dickie et al., 2017), possibilities (1) and (2) seem unlikely for some soaring birds, such as eagles. During spring migration, eagles were on average nearly twice as close to roads and railroads while engaged in slower-paced, stopover-like movements as compared to faster-paced, migratory movements (Fig. 6). We would expect the reverse pattern (i.e. closer association with linear features during more rapid movement phases) if linear features accelerated eagle movement or if eagles were following them during migration.

Slower-paced, tortuous movements are less autocorrelated, and more frequently associated with area restricted search and/or foraging behavior (*sensu*, e.g., Breed et al., 2009; Patterson et al., 2017; Jonsen et al., 2019), so linear features may be targeted as a source of food energy for migrating eagles or serve as open pathways for foraging searches during migratory stopover. Carrion, which is often available along linear features due to vehicle collisions, can be an important food resource for eagles and other raptors (Watson, 2010; Newton, 1979; Bildstein, 2006), and even detection rates of carrion by raptors can be higher near linear features (Lambertucci et al., 2009). Moreover, using linear features to access food aligns with findings in terrestrial mammals; wolves are thought to use seismic lines to access caribou (Dickie et al., 2017), and bears use railroads to access key supplemental food (Murray et al., 2017).

#### Latitudinal patterns in linear feature use

Eagles’ selection for areas close to roads tended to increase with latitude (Fig. 5 & 9), such that selection for roads became stronger as roads became less dense on the landscape, perhaps also coinciding with reduced prey availability. The lower elevation areas eagles used and selected for during spring migration (Fig. 4 & 5) are predominantly boreal forest at high latitudes. Roads and railroads, therefore, contrast markedly with the thick forested landscape, potentially providing open habitat and offering visibility advantages for capturing prey and/or detecting carrion. Other features associated with roads and railroads, such as power poles, likely also offer convenient, high-visibility perches (Meunier et al., 2000).

#### Seasonal patterns in linear feature use

Carrion is usually an ephemeral resource on the landscape; however, it can be more predictable along linear features due to vehicle collisions with wildlife, offering scavenging migrants an efficient and reliable source of food (Lambertucci et al., 2009). While we found eagles exhibit a general preference for areas near roads during both spring and fall migration (Fig. 5), we also found less variable (Fig. 5 & 9) and heavier (Fig. 6–9) use of roadways during spring. With the likely association between roads and foraging, such narrow individual-level variability in selection for roads (Fig. 5 & 9) might be expected from theory, as time- and energy-minimization strategies for migration are tied strongly to fuel deposition rate, leading to selective pressure on en-route foraging strategy (Hedenstrom and Alerstam, 1997). Furthermore, shorter overall migration times in spring than fall—an apparent >50% difference in our sample of eagles—have been found to be owed to differences in foraging-related factors across a number of studies (Nilsson et al., 2013).

Carrion along linear features could be important for an eagle’s timely arrival on the breeding grounds in spring, such that the reward of carrion or benefit of using linear features to detect prey or access carrion could outweigh the real or perceived risk of being struck by a vehicle. By contrast, in fall, there is less urgency for timely arrival on the overwintering grounds, so the elevated mortality risk associated with food resources along linear features would not outweigh the risk. Migrants should, then, spend more time searching for food away from linear features in the fall, which is consistent with our results (Fig. 1 & 6). A key aspect to this reasoning, however, is that individuals assess the mortality risk at linear features correctly. In cases where using a linear feature is more dangerous than an individual’s assessment, use of that linear feature may be maladaptive. For example, high-traffic roads may be easy to assess correctly; however, rural highways with infrequent but fast-moving vehicles may be difficult to assess but quite dangerous.

During fall migration, eagles tended to associate less with linear features as compared to spring (Fig. 6 & 8). This could be a result of more intense pressure to reach summer breeding grounds, but these patterns may simply emerge from eagles responding to relative seasonal abundances of food available across the landscape. In spring, golden eagles migrate earlier than other avian migrants and prior to the emergence of hibernating mammals. In contrast, eagles migrate in fall when migratory birds and mammals not yet hibernating, including many young of the year, are available as prey. Thus, prey may be available across the landscape away from linear features and more easily captured during fall migration compared to spring. Additionally, fall migration coincides with many hunting seasons in western North America, so eagles might also be feeding on unsalvaged large mammal remains (e.g., gut piles) left in the field by hunters away from major roads. Annual moose harvest in British Columbia, Canada averages ∼ 10,000 animals between 15 August and 10 December (Kuzyk, 2016), leaving potentially abundant scavenging opportunities on the landscape. Even if roads provide the same access to carrion during both seasons, in fall the higher relative abundance of prey elsewhere on the landscape would decrease the relative value of roads, which might explain eagles’ less frequent use of linear features in fall. Scavenging opportunities along roads are likely more frequent in late winter and spring, though, due to, for example, accumulated carcasses melting out of snow (Jennelle et al., 2009; Santos and Carvalho, 2011).

### Effects of habitat on eagle space use

Although investigating the effects of anthropogenic linear features was the primary goal of this paper, incorporating other biologically important variables revealed some noteworthy patterns. Unsurprisingly, eagles showed a very strong preference for south-facing slopes (Fig. 5). Given that eagles’ use of south-facing slopes aligned with migratory movements each day (Fig. 4), actively seeking out south-facing slopes is a likely mechanism for how thermal uplift emerges as an important driver of golden eagle movement and behavior during migration (Fig. 3; Eisaguirre et al., 2019; Miller et al., 2016; Duerr et al., 2012). Although biologically expected, this finding provides additional confidence that other estimated parameters are also biologically meaningful.

Although we expected eagles would forego use of south-facing slopes to make more direct routes when under the greater time pressure of spring migration, preference for south-facing slopes actually increased with latitude during spring (Fig. 5 & 9), suggesting eagles balance time- and energy-minimization migratory strategies (Miller et al., 2016); this is also consistent with findings supporting the efficiency of thermal soaring (Duerr et al., 2012). As eagles approach the breeding grounds in spring, thermal uplift and the energetic subsidy it provides are likely more limited due to snow cover increasing with latitude (Eisaguirre et al., 2019), eliciting the greater selective response for south-facing slopes.

Similar to use of linear features, we also found that use of certain habitats was behavior-specific. Use of vegetated areas appeared associated with fly-and-forage movements, identified by lower and intermediate estimates of *γ*_*i*_, and less associated with migratory movement (Fig. 4). In fly-and-forage movement strategies, migrants opportunistically forage while making progress, albeit somewhat more slowly than in directed movement, along the migration route (Strandberg and Alerstam, 2007; Klaassen et al., 2008; Alerstam, 2011). Making wider searches over less than ideal hunting habitat (i.e. a fly-and-forage strategy) might offer occasional food payoffs that, combined with the migration progress, outweigh targeting better hunting areas with more intensive search, which yields no migration progress (Nilsson et al., 2013).

The latter episodic intensive search strategy aligns with the traditional notion of stopover (Gill, 2007); however, migratory “pacing” was recently suggested as a better conceptual framework for stopover behavior in soaring species, such as eagles, that naturally encompasses the fly-and-forage strategy (Eisaguirre et al., 2019). During fall migration when making fly-and-forage movements, eagles used south facing slopes (Fig. 4), perhaps to use thermal uplift to minimize energy expenditure. These periods also coincided with using slightly more vegetated and lower elevation areas (Fig. 4). Use of areas close to waterways also correlated with fly-and-forage movements (Fig. 6), and as many of the waterways encountered along migration routes would be frozen and snow covered during spring migration, these areas could offer open and edge habitat for foraging.

### Individual heterogeneity in movement and space use

There was marked individual-level variability in both behavioral responses to flight subsidies and preference for habitat features (Figs. 3, 5, & 9), which did not seem to be explained by sex (Figs. 3 & S2). In particular, although generally individuals selected areas closer to linear features, some individuals tended to avoid them (Figs. 5 & 9). Similar variation in use of anthropogenic linear features has been reported in grizzly bears *Ursus arctos* (Murray et al., 2017).

Additionally, individual animals can exhibit different habitat preferences depending upon the predominant habitats available to them (Gilbert et al., 2017); however, we did not find such correlations between availability and preference prevalent for individual eagles (Figs. S3 & S4). An exception was for elevation, which could be owed to longitudinal trends in available elevations along the migration routes (Figs. S3 & S5).

Lastly, the effects of learning and variation in personality traits should not be discounted as possible causes for the differences among individuals we found (Fagan et al., 2013; Wolf et al., 2007; Dingemanse et al., 2010). Variation in use of risky sources of food near roads has been shown to exist among an assemblage of raptor species that exhibit a range of risk aversion behavior (Lambertucci et al., 2009). This variation should also exist among individuals within species; each individual would have a unique risk-aversion personality (Wolf et al., 2007; Dingemanse et al., 2010). Thus, it is reasonable to expect that some of the variance among eagles in our sample, in terms of their attraction/repulsion to linear features (Figs. 5 & 9), could be due to variance in personality along the shy-bold spectrum and/or behavioral conditioning.

### Conclusions & Implications

One inference drawn from our results reflects how time of day and thermal uplift relates to eagles’ use of linear features. This raises the question of how uplift drives migrant abundance along linear features and whether or not poor uplift conditions might elevate vehicle collision risk for migrants. While we have shown that eagle behavior correlates with use of areas close to roads and railroads and that they can prefer such areas, we could not infer mortality risk. Limited thermal uplift and how it effects raptor behavior has been suggested to elevate risk of raptor collisions with wind turbines though (Barrios and Rodríguez, 2004). Higher frequency telemetry and observational studies along roads during migration could help reveal and perhaps quantify vehicle collision risk; a network of carrion baited camera traps might also be useful and cost effective (*sensu* Jachowski et al., 2015). The parameterized el-SSF, however, could be used to predict eagle space use patterns to identify potential hot spots of elevated mortality risk, as well as how those spots might change with weather. Such application could at least increase the efficiency and effectiveness of targeted studies of mortality risk within this migration corridor, which spans half the continent (Fig. 1), as well as other expansive movement corridors. Moreover, the el-SSF could also be used to help inform eagle-vehicle collision risk models (e.g., Lonsdorf et al., 2018), and it could more generally be used to predict and test hypotheses regarding changes in eagle space use patterns following continued development of linear futures, other habitat changes, and/or changes in weather patterns.

As continued human development has the potential for introducing additional mortality risk onto the landscape, it is important to keep in mind that anthropogenic mortality is considered the greatest threat to many populations of long-lived raptors (Newton, 2008). Further, elevated mortality during migration—an already risky time for birds— can carry over to impact population reproductive rates (Newton, 2008; Harrison et al., 2011). Such carry-over effects were suggested as a possible cause for the long term decline in reproductive success of long-distance migratory golden eagles in a study area in interior Alaska (McIntyre and Schmidt, 2012). While there are apparently few wind energy developments along the migration corridors of our sample of eagles (Fig. S1), there will certainly be continued development, potentially creating more opportunities for migratory golden eagles to interact with wind turbines. Here, we have clearly shown the importance of considering the effects of linear features on avian movement and space use during migration. We thus should not discount the potential movement-related effects and mortality risk that anthropogenic linear features can impose on avian migrants, despite a bird’s ability to fly over such human infrastructure during migration.

## Supporting information

Example data

Stan program

## Author contributions

All authors conceived the ideas of this research and collected the data. JME analyzed the data and led the manuscript. All authors contributed to drafts of the manuscript.

## Acknowledgements

T. & D. Hawkins, M. Kohan, B. Robinson, and many others provided support in the field, and J. Liguori and N. Paprocki helped age eagles. M. Auger-Mèthè offered motivating and invaluable discussion regarding the movement model. C. McIntyre, K. Kielland, and P. Doak provided excellent feedback that helped improve this research and manuscript. To all of these friends, we are most grateful. Funding was provided by the Alaska Department of Fish & Game (ADF&G) through the federal State Wildlife Grant Program, and the U.S. Fish & Wildlife Service (USFWS) provided additional PTTs and data. JME was supported by the Calvin J. Lensink Fund during part of the project. The findings and conclusions of this paper are those of the authors and do not necessarily represent the views of the USFWS.

## Data accessibility

All movement data used for this manuscript are archived in the online repository Move-bank (https://www.movebank.org/; IDs 17680093 and 19389828). The data contain information considered confidential and sensitive by the State of Alaska (State Statute 16.05.815(d)), but they could be made available for research at the discretion of the Alaska Department of Fish & Game and U.S. Fish & Wildlife Service. Code to fit the movement model to data and sample from the posterior predictive distribution is provided as supporting information.

## Supporting Information

## Appendix S1: wind energy developments

Locations of commercial grade wind turbines were gathered from the United States Geological Survey US Wind Turbine Database, EnergyBC, and Alberta Human Footprint Monitoring Program. Coordinates for the two wind turbines in the Yukon Territory were approximated using Google Earth. Only those turbines indicated as operational or were projected to be operational during each year of the eagle migrations were included when distances to turbines and buffer intersections were computed.

Only 25 of the 17,386 steps analyzed intersected 1 km buffers surrounding wind turbines. Twenty endpoints of the steps were within 5 km of a wind turbine, and only two were within 2 km. Of those 45 cases (of either step intersection or endpoint within 5 km), only four occurred in the USA, with the remainder in Canada, and four individuals were responsible for 33 of them. These findings indicate that areas containing wind energy developments are likely not available at high frequency to the population of eagles we have sampled—golden eagles that summer in southcentral Alaska—during migration, but perhaps isolated to a few locations used repeatedly by certain individuals. We thus chose not to include effects of wind energy developments in candidate models of selection, as the ability to detect a true effect on the scale of inference possible with the data on hand would be extremely low. It is important to keep in mind that there are currently relatively few wind energy developments along the routes of these golden eagles, so further developments could increase the availability of them to Alaska’s migratory golden eagles.

**Figure S1:**
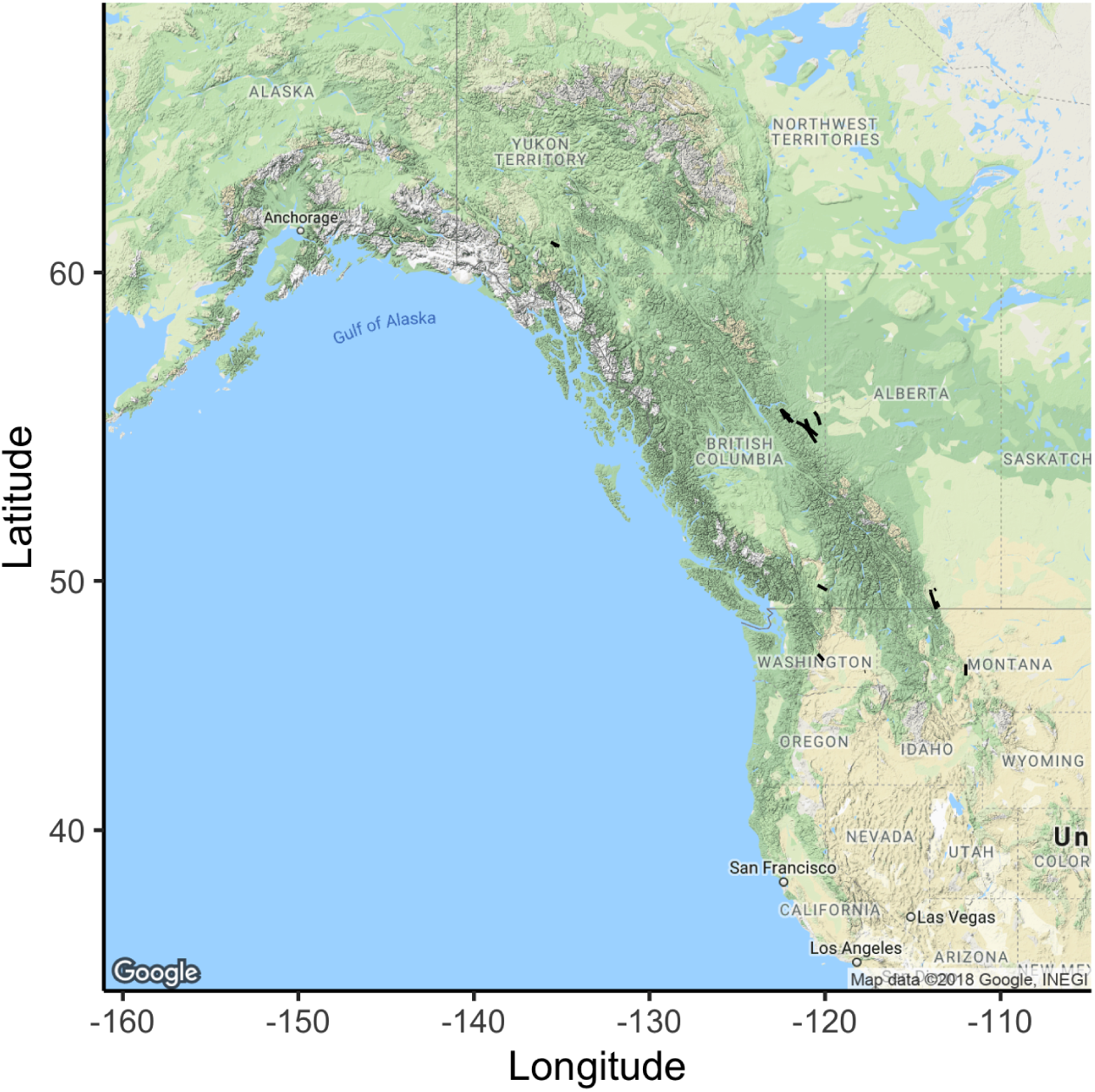
Golden eagle migration track segments that intersect a 1-km buffer around a wind turbine or with a point that is within a 5 km of a wind turbine.

## Appendix S2: supplementary tables & figures

**Figure S2:**
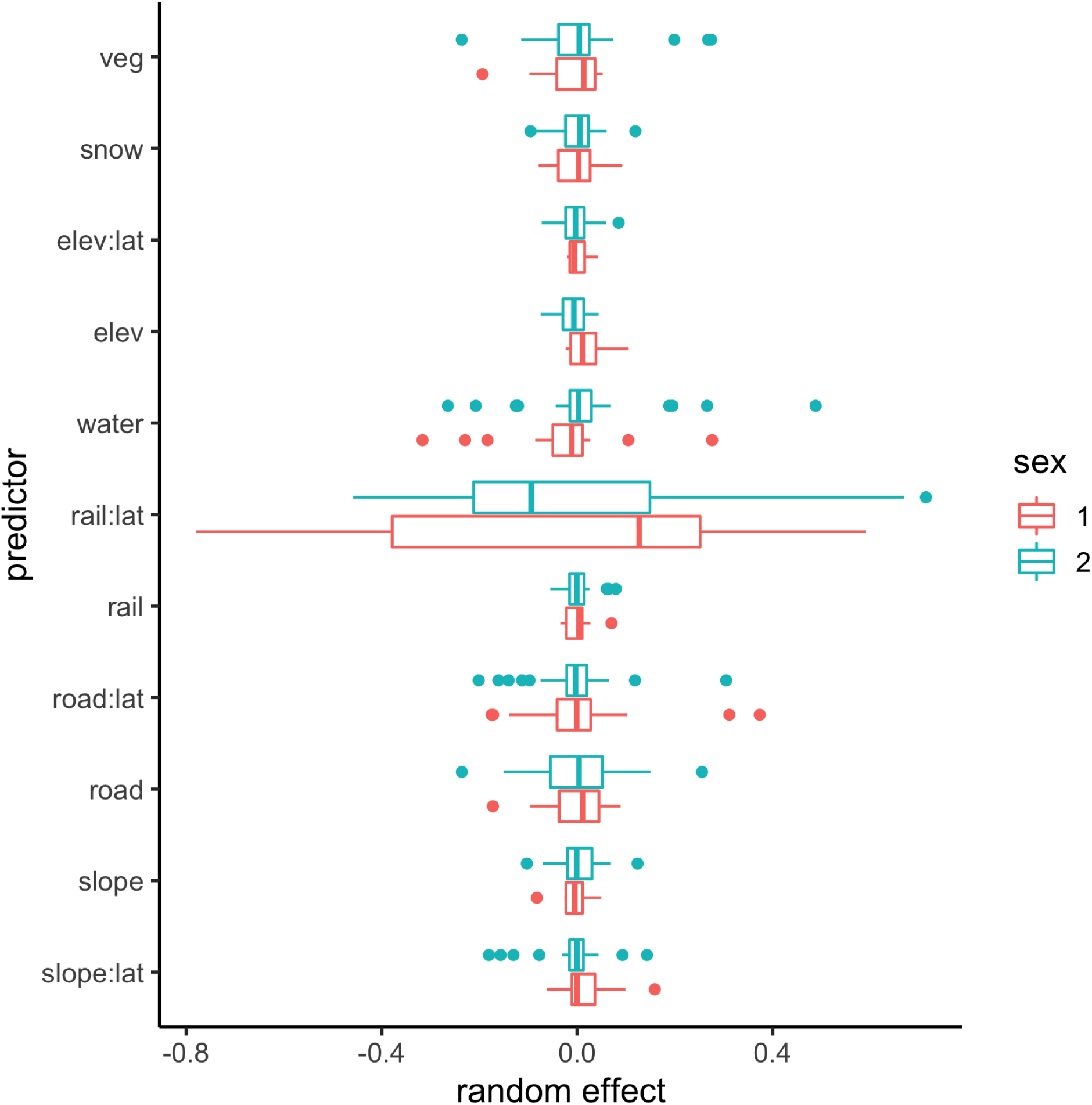
Individual-level golden eagle migration habitat selection effects estimated with a step selection function shown grouped by sex.

**Figure S3:**
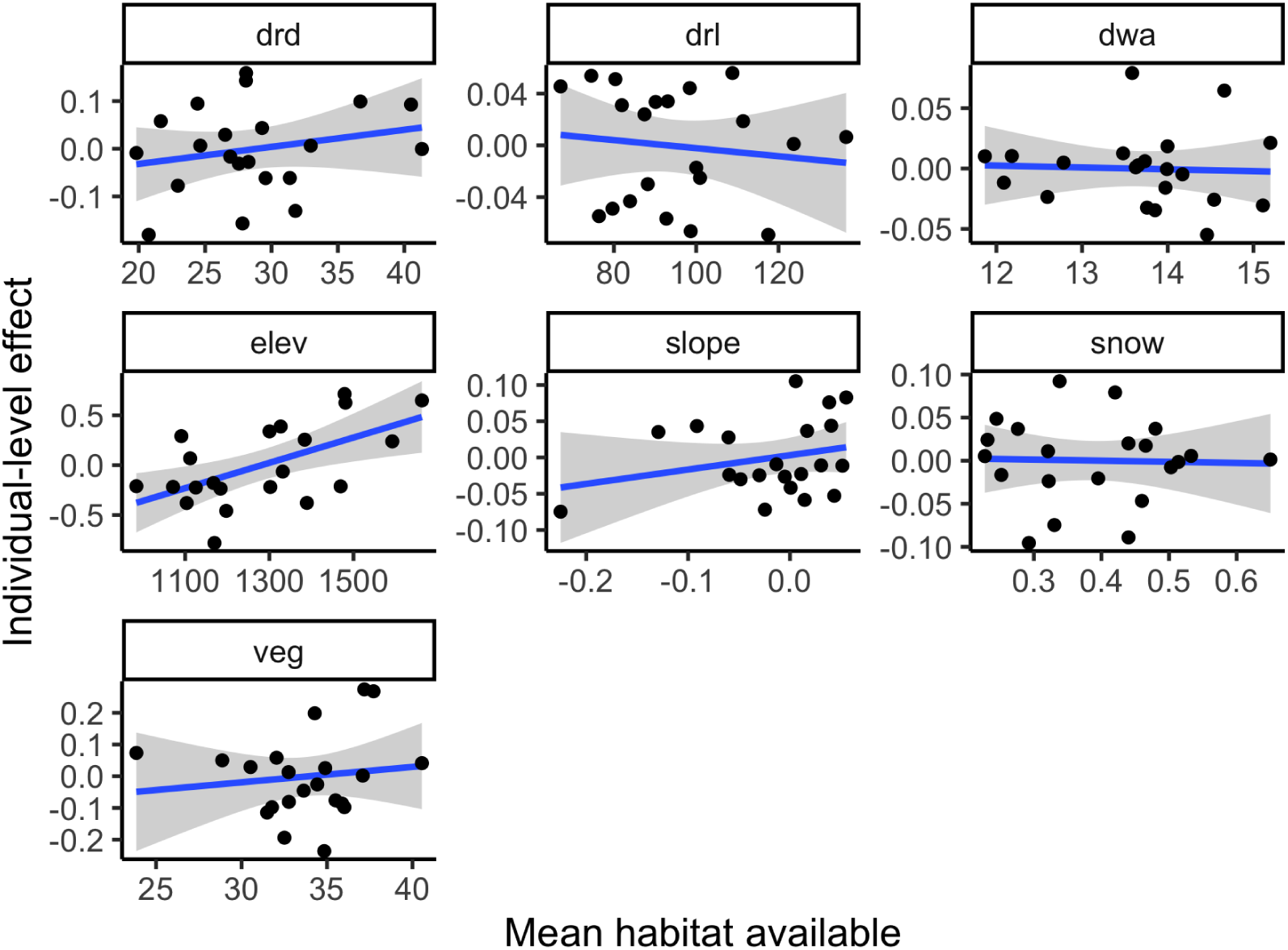
Effects of habitat available during fall migration on individual-level habitat selection effects. ‘drd’, ‘drl’, ‘dwa’, ‘elev’, ‘slope’, ‘snow’, and ‘veg’, correspond to distance to nearest road, distance to nearest railway, distance to nearest waterway, elevation, slope southing, percent snow cover, and percent vegetation cover, respectively. Spearman’s rank correlation tests indicated only available elevation correlated significantly with selection for elevation (*p* < 0.05). Shaded areas represent 95% confidence intervals of the simple linear regressions.

**Figure S4:**
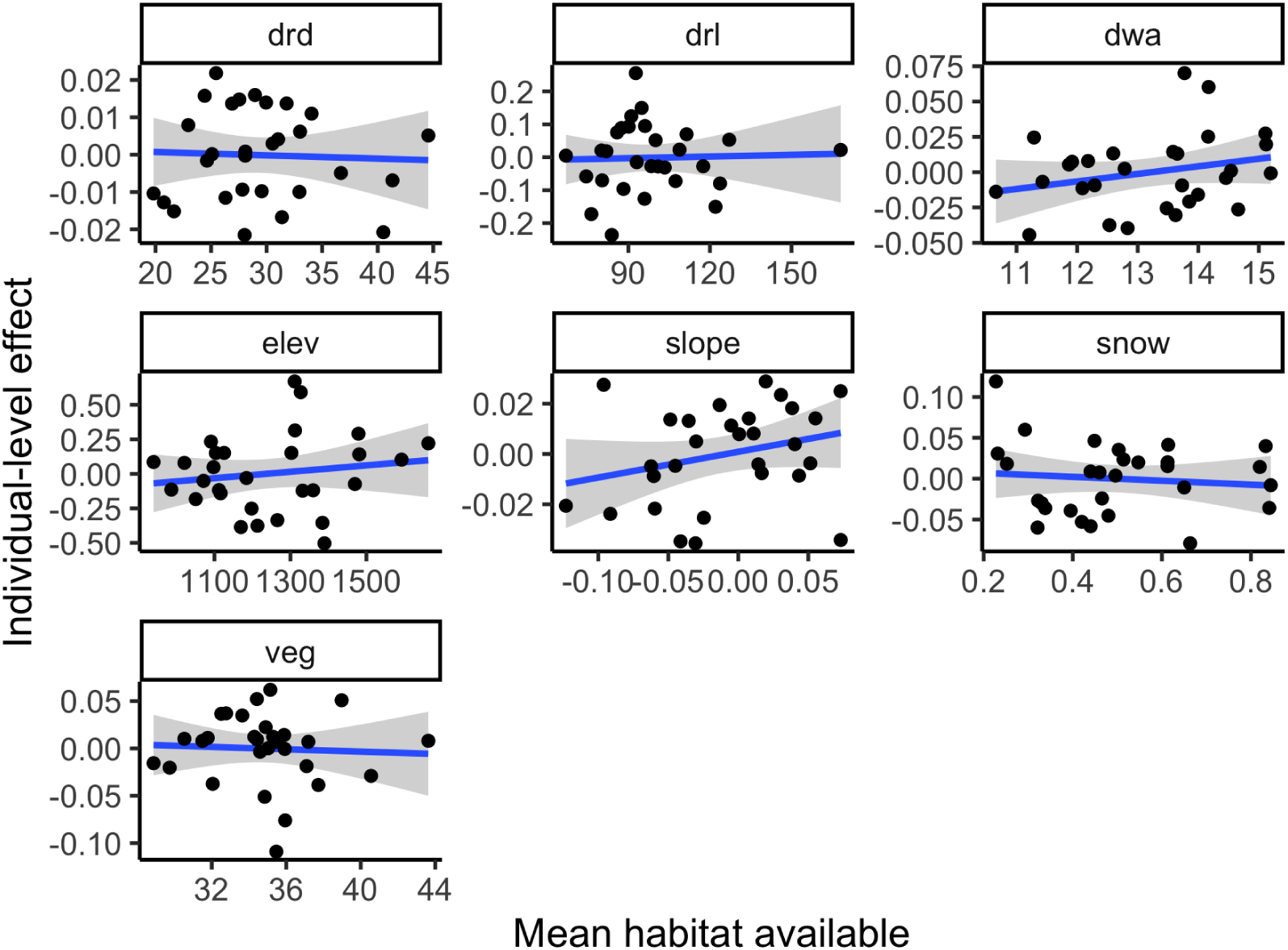
Effects of habitat available during spring migration on individual-level habitat selection effects. ‘drd’, ‘drl’, ‘dwa’, ‘elev’, ‘slope’, ‘snow’, and ‘veg’, correspond to distance to nearest road, distance to nearest railway, distance to nearest waterway, elevation, slope southing, percent snow cover, and percent vegetation cover, respectively. None of the correlations were statistically significant. Shaded areas represent 95% confidence intervals of the simple linear regressions.

**Figure S5:**
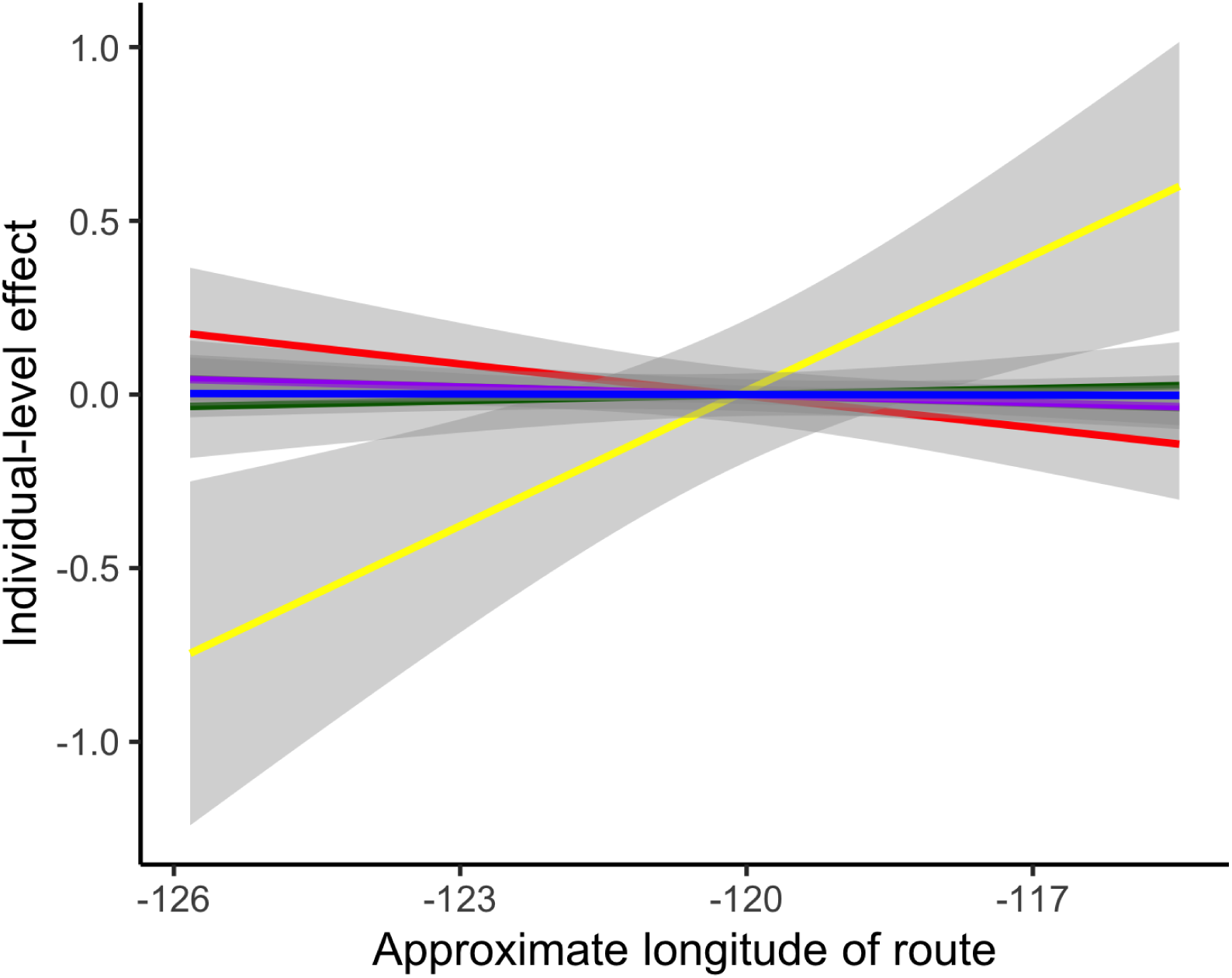
Effects of longitudinal location of individual eagles’ routes (when crossing 53°N latitude) on individual-level habitat selection effects during fall migration. Yellow curve corresponds to elevation. Shaded areas represent 95% confidence intervals of the simple linear regressions.

**Figure S6:**
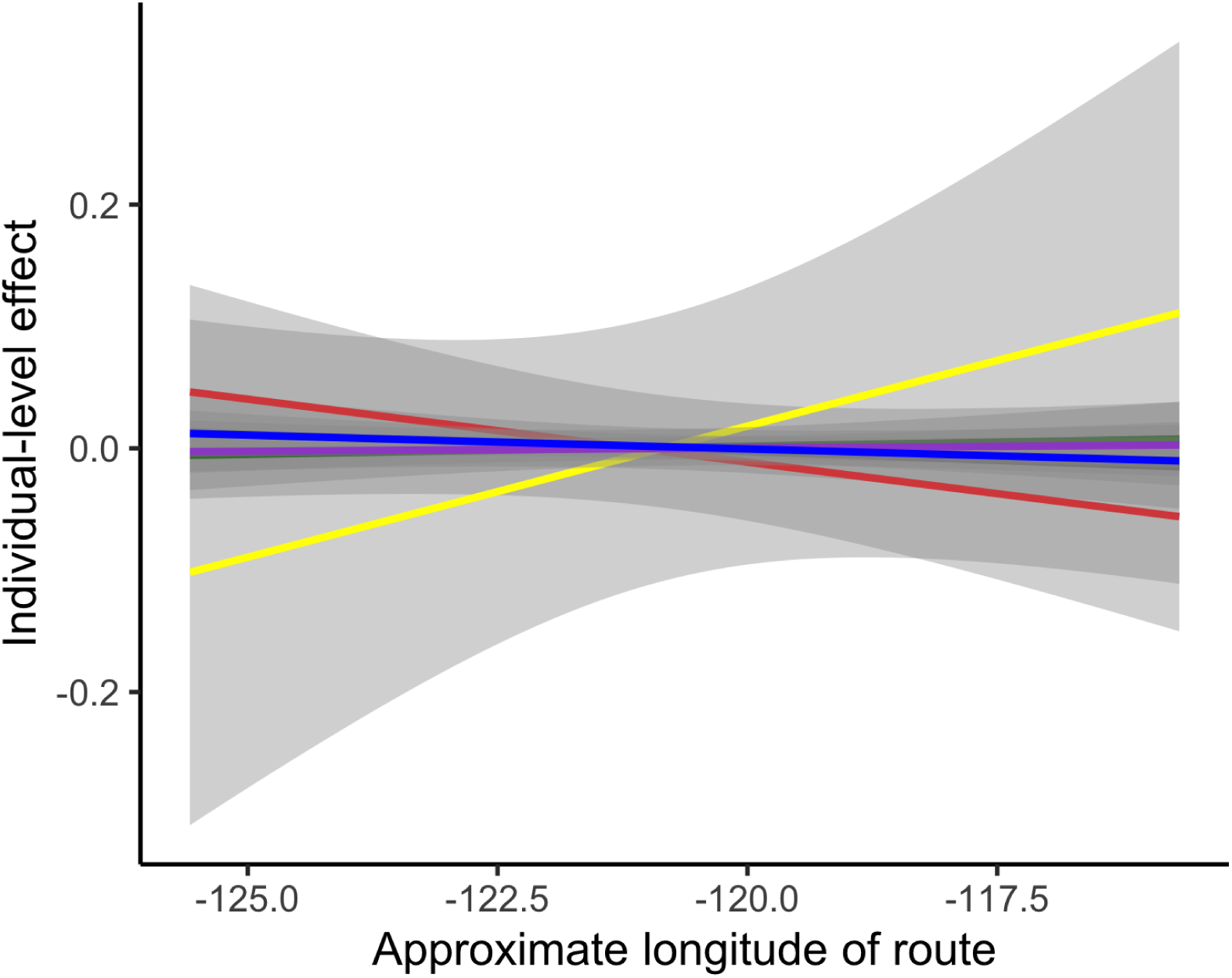
Effects of longitudinal location of individual eagles’ routes (when crossing 53°N latitude) on individual-level habitat selection effects during spring migration. Yellow curve corresponds to elevation. Shaded areas represent 95% confidence intervals of the simple linear regressions.

**Figure S7:**
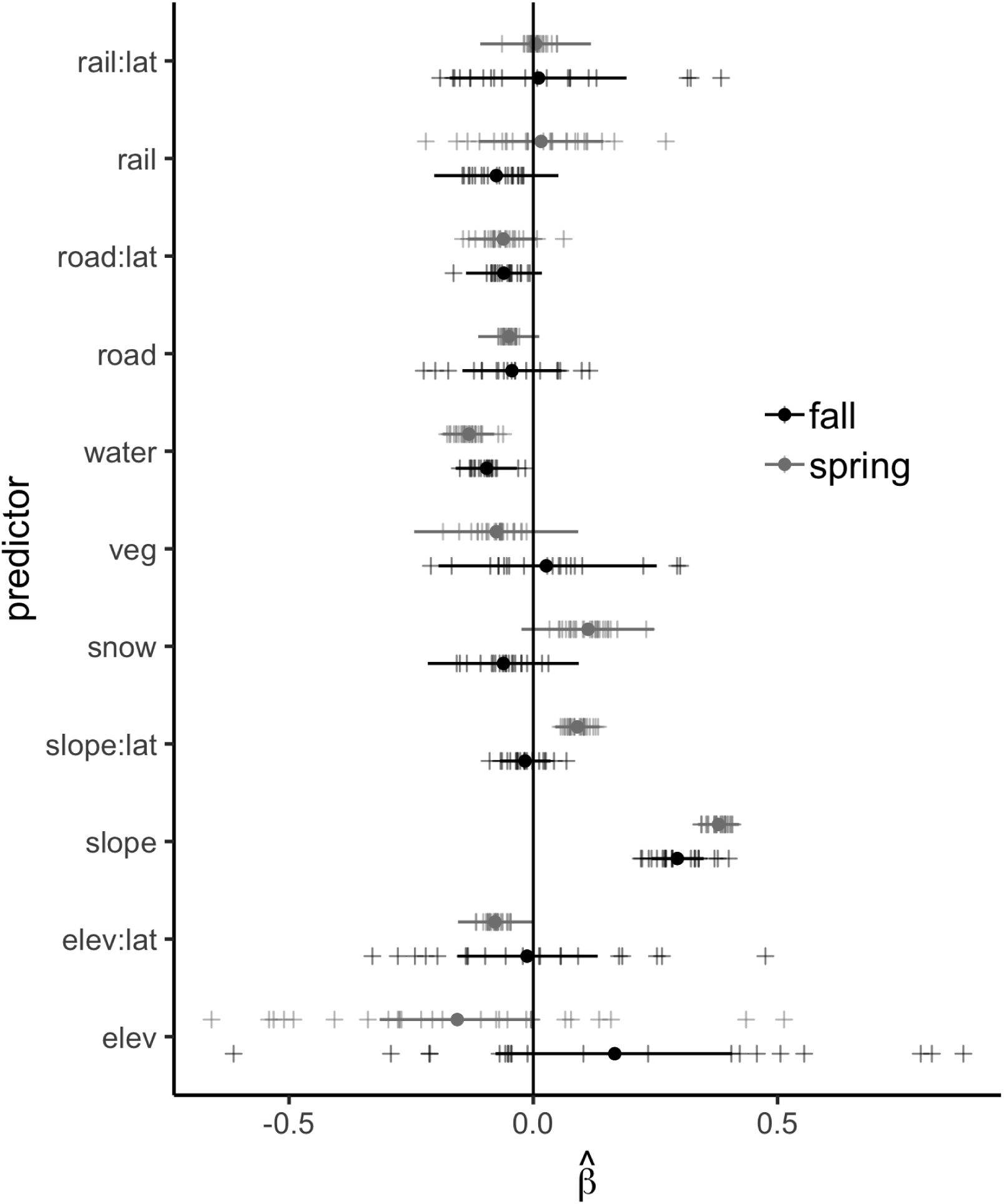
Full version of figure 5 in the main text.

## Appendix S3: simulation study

The goal of this brief simulation study was to ensure that a step selection function (SSF) similar to the one used for the study presented in the main text can recover a known effect a linear feature on animal space use, as well as discern occasions when a linear feature does not affect space use. Over real landscapes, some linear features often occur parallel to each other (e.g., roads and railroads, roads and transmission lines, etc.), and many animals might move in parallel to linear features if, for example, their migration corridors are aligned with those features. We were, thus, primarily interested in assuring the SSF yields robust inferences under these scenarios, both when a linear feature does and does not affect space use.

To test these points, we simulated 10,000 tracks of 100 steps in length over a hypothetical landscape void of features except for two straight, parallel linear features (Fig. S8). The tracks were simulated from the movement model presented in the main text (equations 3-4) modified slightly with a directional bias paralleling the linear feature plus *γ*_*i*_ modeled as a sine wave. We approximated the fitted movement kernel with a bivariate Gaussian kernel and imparted additional random noise on *γ*_*i*_ to reflect imperfect fit of the movement kernel. The conditional logistic regression was then carried out for each simulated track with five available points drawn from the the ‘fitted’ movement kernel using `clogit ()` from the `survival` package in R.

In the simulations, selection for space nearer the linear feature of interest was imparted by shifting each point in each track 10% closer to that linear feature, while leaving the availability (i.e. the ‘fitted’ movement kernel) unchanged. For each simulated track and respective fit, we retained the estimated coefficients 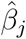 for distance to each linear feature, where *j* = 1 for the linear feature selected for and *j* = 2 for the parallel, and the resulting *p*-values from the tests of *H*_0_: 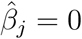.

We found that the SSF performed well in recovering selection for the linear feature, and additionally it successfully showed when a parallel linear feature was not selected for (Fig. S9). As expected, the SSF did not find (at *α* = 0.05) 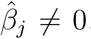, *j* = 1,2 for 94% percent of the simulated tracks prior to selection for the linear feature being imparted. After selection was imparted on each track, the SSF found 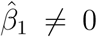 for 91% of the tracks and 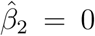 for 72%. More explicitly, the SSF correctly identified when both linear features were not selected for in 94% of the simulations; it correctly identified the selection in 91% of the simulations; and it correctly identified no selection for the parallel linear feature in 72% of the simulations when selection for the other linear feature was imparted. While it was not correct (based on *p*-values) in 29% of the simulations when there was no selection for the parallel feature, this simulation study represents a sort of worst case scenario, where two linear features are perfectly parallel over a large portion of a study area. We would thus expect even better performance of the SSF over a real landscape, where such perfect correlation would occur much less frequently.

**Figure S8:**
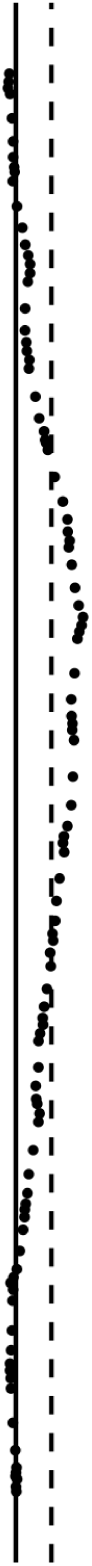
One simulated track over the hypothetical landscape with two parallel linear features. In the simulation study, selection was imparted for the solid line, and the dashed line was the parallel linear feature.

**Figure S9:**
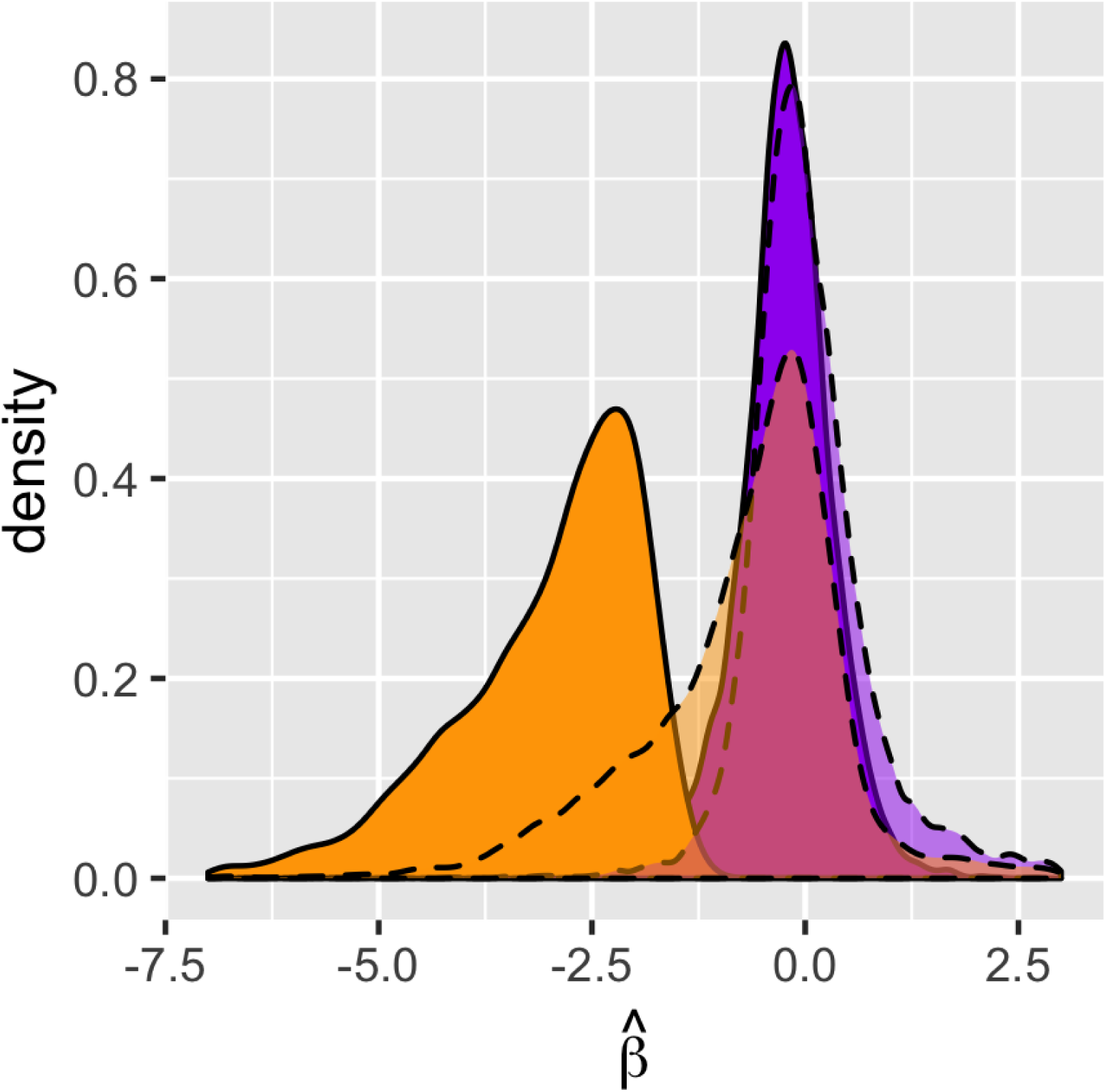
Density plots of estimated selection coefficients from the simulation study. Dashed curves are the densities of the estimated selection coefficients for the parallel linear feature. Orange densities correspond to estimated selection for the linear features after selection was artificially imparted. Purple densities are of the estimates with no selection imparted.

