## Supplementary material for "Novel step selection analyses on energy landscapes reveal how linear features alter migrations of soaring birds": R code

library(rstan)
library(loo)

#################################
## model variables
# x -- vector of x coordinates
# y -- vector of y coordinates
# dt -- vector of time intervals
# N -- number of fixes in track
# oro -- vector of raw orographic uplift data
# therm -- vector of raw thermal uplift data
# twind -- vector of raw tailwind data
# tod -- vector of times of day (1=day, 0=night)
################################

dat = read.csv('examp_dat.csv')  

x = dat$x
y = dat$y
dt = dat$dt
N = nrow(dat)
oro = dat$oro
therm = dat$therm
twind = dat$twind
tod = dat$tod

# fit Stan model with HMC, using default no-u-turn sampler
# model code should be saved as a .stan file
# arguments should be modified as required
stan.fit = stan("ssf_move_code.stan", 
                data = list(x, y, dt, N, oro, therm, twind, tod), 
                pars = c('gamma', 'sigmax', 'sigmay', 'sigmav', 'beta', 'x_new', 'y_new'),
                chains = 2, 
                iter = 1000,
                thin=1,
                cores = 1,
                control = list(adapt_delta = 0.9)) 
# last argument helps with divergent transitions


# draw 5 available points for the ith used point (can loop over i)
i=1
x.av = sample(unlist(rstan::extract(stan.fit, pars = paste0('x_new[',i,']')), use.names=F), 5)
y.av = sample(unlist(rstan::extract(stan.fit, pars = paste0('y_new[',i,']')), use.names=F), 5)
